# Megabodies expand the nanobody toolkit for protein structure determination by single-particle cryo-EM

**DOI:** 10.1101/812230

**Authors:** Tomasz Uchański, Simonas Masiulis, Baptiste Fischer, Valentina Kalichuk, Alexandre Wohlkönig, Thomas Zögg, Han Remaut, Wim Vranken, A. Radu Aricescu, Els Pardon, Jan Steyaert

## Abstract

Nanobodies (Nbs) are popular and versatile tools for structural biology because they have a compact single immunoglobulin domain organization. Nbs bind their target proteins with high affinities while reducing their conformational heterogeneity, and they stabilize multi-protein complexes. Here we demonstrate that engineered Nbs can also help overcome two major obstacles that limit the resolution of single-particle cryo-EM reconstructions: particle size and preferential orientation at the water-air interface. We have developed and characterised novel constructs, termed megabodies, by grafting Nbs into selected protein scaffolds to increase their molecular weight while retaining the full antigen binding specificity and affinity. We show that the megabody design principles are applicable to different scaffold proteins and recognition domains of compatible geometries and are amenable for efficient selection from yeast display libraries. Moreover, we used a megabody to solve the 2.5 Å resolution cryo-EM structure of a membrane protein that suffers from severe preferential orientation, the human GABA_A_ β3 homopentameric receptor bound to its small-molecule agonist histamine.

## INTRODUCTION

Single particle cryo-EM has recently become the technique of choice for the structural characterisation of membrane proteins and macromolecular complexes^1–3^. Instrumentation and data analysis methods continue to improve at a spectacular pace^4,5^. However, factors including small particle size, low symmetry, high flexibility and non-random distribution in ice limit the achievable resolution of 3D reconstructions. Whereas large molecules are relatively easy to identify in noisy low-dose images of frozen hydrated samples, and have sufficient features to facilitate the accurate determination of their orientation and position required for alignment and averaging^6^, structural analysis of small particles (∼100 kDa or less) is considerably more difficult. Phase contrast methods such as defocusing the objective lens or the use of phase plates, especially the Volta phase plate (VPP)^7^, enable imaging of biological specimens with an acceptable (albeit still limiting) amount of radiation damage^8^. Despite the recent success in high-resolution reconstructions of small macromolecule proteins aided by a VPP including streptavidin (52 kDa tetramer, ∼3.2 Å)^9^ and human haemoglobin (64 kDa tetramer, ∼3.2 Å)^10^ or using conventional defocus-based data collection for methaemoglobin (64 kDa tetramer, ∼2.8 Å and ∼3.2 Å) and alcohol dehydrogenase (82 kDa dimer, ∼2.7 Å)^11^, the routine analysis of such samples remains very challenging. To circumvent this problem, several protein engineering approaches have been employed to increase the size of particles. Small proteins have been genetically fused to scaffold proteins in order to increase their size directly^12^ or bound to recognition domains organised in a large molecular support of high symmetry^13,14^. Attempts to align the target molecules according to the symmetry of a multimeric scaffolds, with the help of local/focused classification techniques, are useful in principle but in practice limited by the flexibility of linker regions.

Regardless of particle size, their distribution in the thin ice layer greatly impacts on the quality and resolution of resulting cryo-EM maps^15^. The idea that particles are randomly oriented in ice after grid blotting and rapid cryo-plunging is far from the experimental reality. During the vitrification process, particles diffuse and interact with the water-air and/or water-support interfaces even 1000 times a second^15,16^. This can lead to protein denaturation but also, in almost all instances, particles are adsorbed to such interfaces and present preferential orientation due to their surface properties^17^. While supplementation of buffers with surfactants such as detergents is often employed to improve particle distribution in ice^18^, for sensitive specimens including membrane proteins this approach should be avoided. To limit the impact of preferential particle orientation, specimens can be tilted inside the microscope during data collection^19^. Additionally, faster plunge-freezing devices that reduce the time between sample application and vitrification^20^, utilization of graphene-supported grids with tuneable surface properties^21^ or engineered hollow 3D-DNA structures able to trap DNA-binding proteins inside the scaffold^22^ are also being explored. Yet, to date, such approaches could not be generalised to eliminate the particle orientation problem in cryo-EM.

To help overcome these performance barriers, we designed a novel class of chimeric molecules, called megabodies (Mbs). Megabodies are built by grafting single domain antigen-binding constructs such as nanobodies (∼15 kDa)^23,24^ or monobodies (∼10 kDa)^25^, into larger scaffold proteins to produce stable, reasonably rigid and efficiently-folding monomeric chimeras. The generic nature of this technology enables a direct selection of novel megabodies directly from nanobody immune libraries or, alternatively, a straightforward reformatting of existing nanobodies. We show that megabodies, unlike their parental nanobodies, successfully overcome the severe preferential orientation of a relatively small (∼200 kDa) membrane protein, the human GABA_A_-β3 receptor reconstituted in a lipid nanodisc. The excellent quality of the cryo-EM map obtained, to 2.5 Å resolution, allows unambiguous identification and refinement of histamine, a small-molecule agonist composed of only eight non-hydrogen atoms.

## RESULTS

### Design, expression and purification of megabodies

The generic megabody design consists of a small globular domain, such as a nanobody, grafted into a scaffold protein via two short peptide linkers (**Fig. 1a, b**). As proofs of principle, we designed synthetic genes encoding megabodies build from a GFP-specific nanobody (Nb207, **Supplementary Table 1**) that is linked through its first loop (connecting β-strands A and B) to an exposed β-turn of selected scaffold proteins. We have identified two secreted bacterial proteins containing antiparallel β-strands with surface accessible β-turns: β-turn S3-S4 of the adhesin domain of *H. pylori*^26^ (HopQ, 45 kDa, PDB ID: 5LP2, **Supplementary Fig. 1a-c**) and β-turn A’S1-A’S2 of the *E. coli* K12 Glucosidase YgjK^27^ (YgjK, 86 kDa, PDB ID: 3W7T) (**Supplementary Fig. 1d-e**). Accordingly, we grafted Nb207 into circular permutants^28^ of these molecular scaffolds to build megabodies 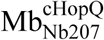 (**Fig. 1c**) and 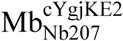 of about 56 and 100 kDa, respectively (**Supplementary Fig. 2**). These chimeric proteins were next produced as highly soluble proteins in the periplasm of *E. coli* and purified in milligram quantities (**Supplementary Fig. 2f, g**). Thermal shift assays indicate that both megabodies are conformationally stable, characterised by melting temperatures (T_m_) of approximately 50 °C (**Supplementary Fig. 2h**). In our experience, these megabodies also resist at least two freeze-thaw cycles.

**Figure 1.**
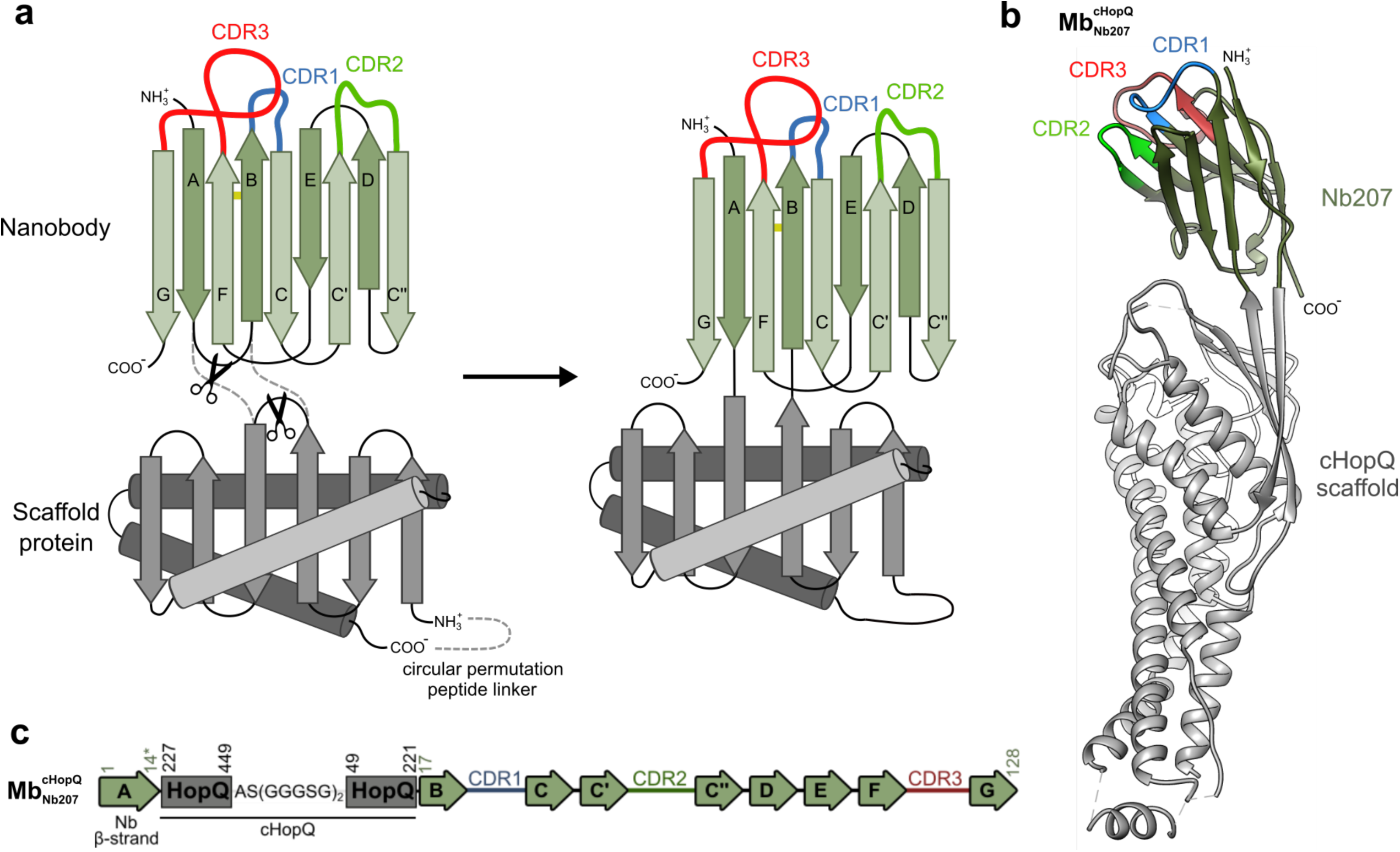
Molecular design of novel rigid antibody chimera called megabodies. **a**, Megabodies are assembled from a nanobody (or a similar single-domain antigen-binding protein) and a (large) scaffold protein. The (optionally circularly permutated) scaffold protein is placed between β-strand A and β-strand B of a nanobody. The megabody is encoded by a single gene, comprising a nanobody that is grafted into the scaffold protein via two peptide bonds. **b**, Crystal structure of 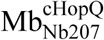 (PDB code: 6QD6) that was built from a GFP-specific nanobody (Nb207) and a circularly permutated variant of HopQ (cHopQ). Molecule A, one out of ten molecules present in the asymmetric unit (RMSD range between 0.557-3.341 Å) is represented. CDRs and β-strands of the nanobody are defined according to IMGT. **c**, Schematic representation of the primary structure of 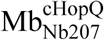. The AS(GGGSG)_2_ peptide was used to circularly permutate HopQ. β-strands are represented by arrows. Residues of the nanobody and HopQ fragments are numbered according to IMGT and UniProtKB B5Z8H1 numbering, respectively.

**Figure 2.**
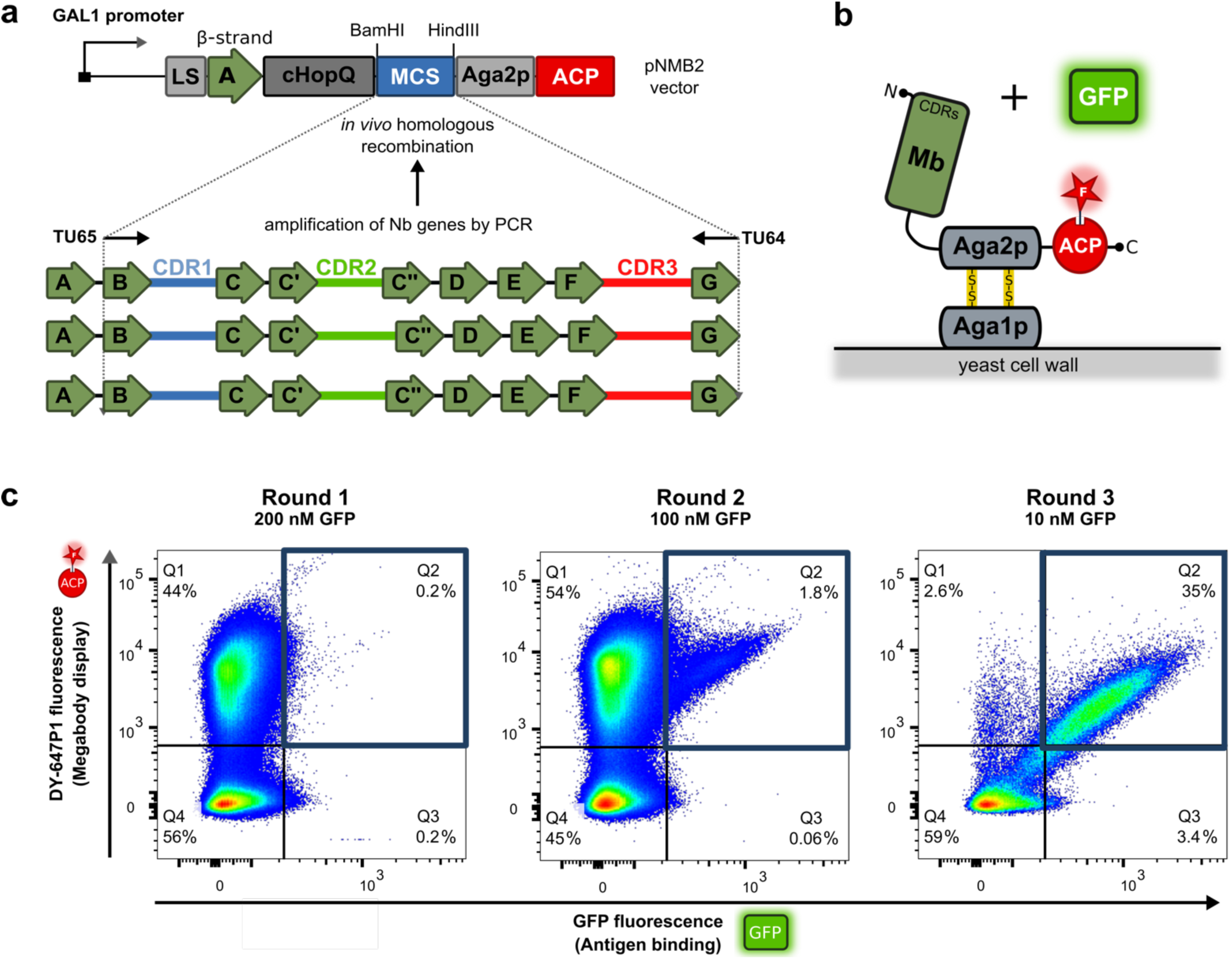
Selection of megabodies from nanobody immune libraries by yeast display. **a**, Gene fragments encoding β-strands B to G of a nanobody immune repertoire can be amplified by PCR using primers TU65 and TU64 to be cloned into yeast vector pNMB2 that encodes a display cassette containing the following elements: the appS4 leader sequence (LS) to direct secretion, a consensus sequence encoding the conserved β-strand A of the nanobody fold (A), a circularly permutated variant of HopQ (cHopQ), a multicloning site (MCS) to clone the nanobody immune library, the Aga2p anchor protein followed by an acyl carrier protein (ACP). **b**, The display level of a cloned Megabody on the surface of a single yeast cell can be monitored through a covalent fluorophore (red star) that is attached in a single enzymatic step to the ACP tag. Antigen binding to the displayed Megabody can be monitored simultaneously by following the fluorescence of the antigen (in this case GFP). **c**, Dot plots representing the enrichment of GFP-specific megabodies within three constitutive rounds of yeast display selection. The y-axis quantifies the megabody display level (DY-647P1 fluorescence), whereas x-axis is a measure of the amount GFP that is bound to the cell. Yeast cells displaying GFP-specific megabodies (gate Q2 represented as a blue square), were enriched from 0.2% in the first round (200 nM of GFP) to 35% in the third round of selection (10 nM of GFP).

To assess the structural impact of the proposed fusion scheme, we grew diffracting crystals of both megabodies. Although attempts to obtain a structure of 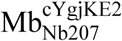 failed so far, partly due to poor diffraction (∼5 Å), we solved the crystal structure of 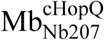 to 2.8 Å resolution (**Fig. 1b, Supplementary Fig. 3, Supplementary Fig. 4, Supplementary Table 2**). The 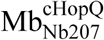 crystallised in the space group P1 containing ten molecules in the asymmetric unit (PDB ID: 6QD6). The two short peptide linkers connecting the nanobody and the scaffold are well-defined in the electron density map, except for molecule J, where the first twelve residues of the megabody were not visible. The ten molecules in the asymmetric unit superimpose with low root mean square deviations (RMSD) in a range from 0.557 to 3.341Å (for 1-532 equivalent C_α_ positions refined in the electron density), mainly caused by the nanobody-scaffold linker flexibility which allows a maximal relative rotation of 22.2° when measured between molecules B and E (**Supplementary Fig. 3d**). The circular permutation and the insertion of the nanobody between β-strands S3 and S4 of HopQ does not affect the overall tertiary structure of this scaffold protein (**Supplementary Fig. 4**). The elongated shape of 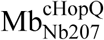 revealed by crystallography is in agreement with small angle X-ray scattering (SAXS) analysis performed in solution, where the crystallography-based theoretical scattering profile yielded overall good fit to the experimental data (**Supplementary Fig. 5**, discrepancy *χ*^2^_SAXS_ in a range from 1.699 to 2.033 for molecule B and E, respectively). Most importantly, Nb207 and both its enlarged variants, 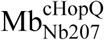 and 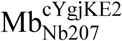 bind their cognate antigen GFP with similar affinities (**Supplementary Fig. 6**), which confirms that the binding properties of the parental nanobody are not affected by insertion into different scaffolds according to **Fig. 1a**.

**Figure 3.**
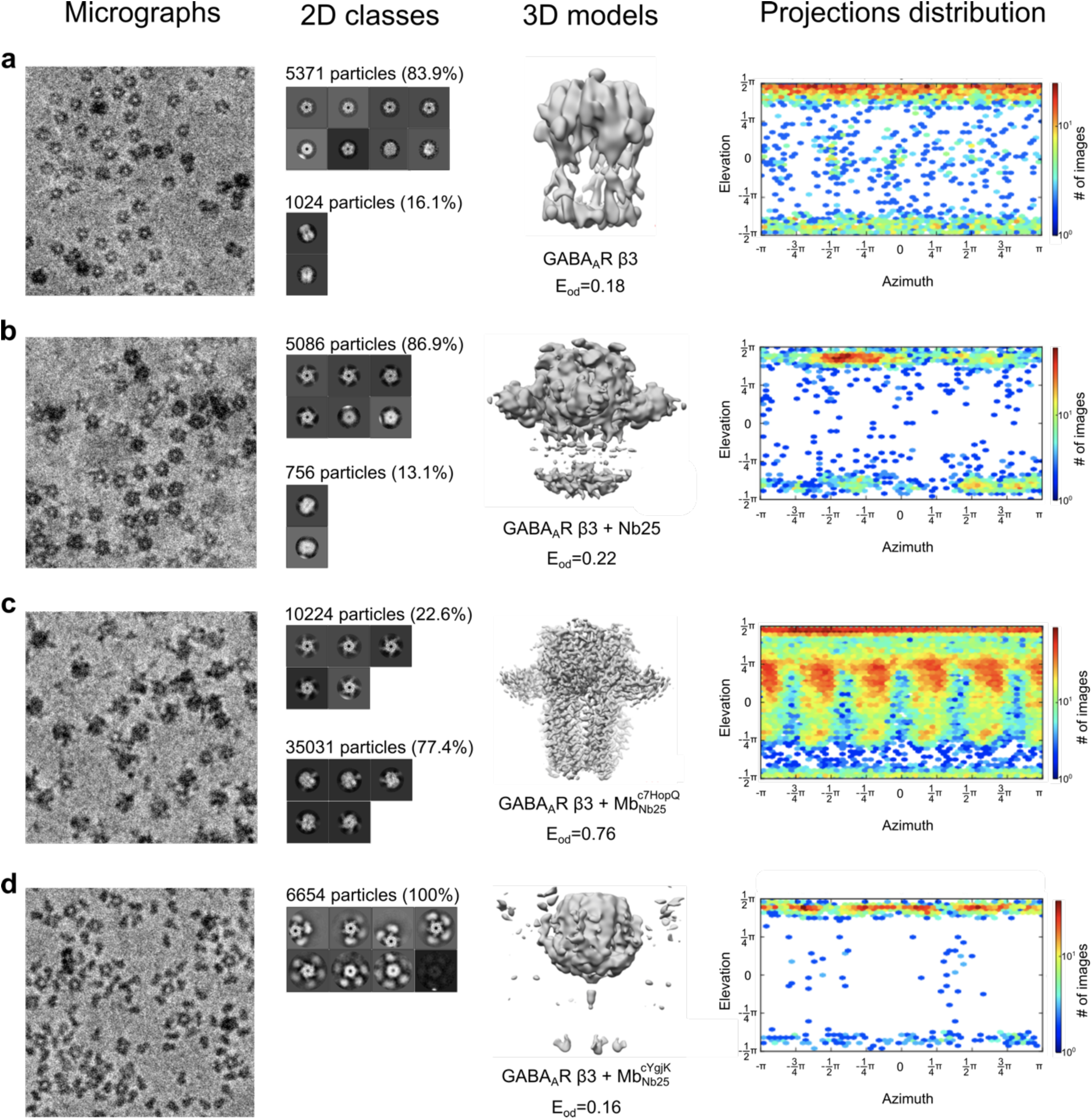
Cryo-EM datasets of the homomeric GABA_A_ β3 receptor alone, in complex with Nb25 or bound to megabodies derived from Nb25. **a-c**, Direct comparison of homomeric GABA_A_ β3 receptor alone (**a**), with addition of Nb25 (**b**), 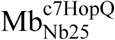 (**c**) and 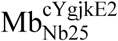 (**d**) in single particle cryo-EM. Aligned representative micrographs and 2D classes for each experimental sample are shown in far and middle left panels, respectively. 2D classes were separated into two groups: top-view (top) and tilted-view (bottom). Counts of particles with a top-view or tilted-view orientation are indicated. Reconstructed 3D models and corresponding particle distribution “efficiencies” (E_od_) for every condition are shown in middle right panels. Distributions of viewing directions over azimuth and elevation angles (far right panels) are calculated for shown 3D models.

**Figure 4.**
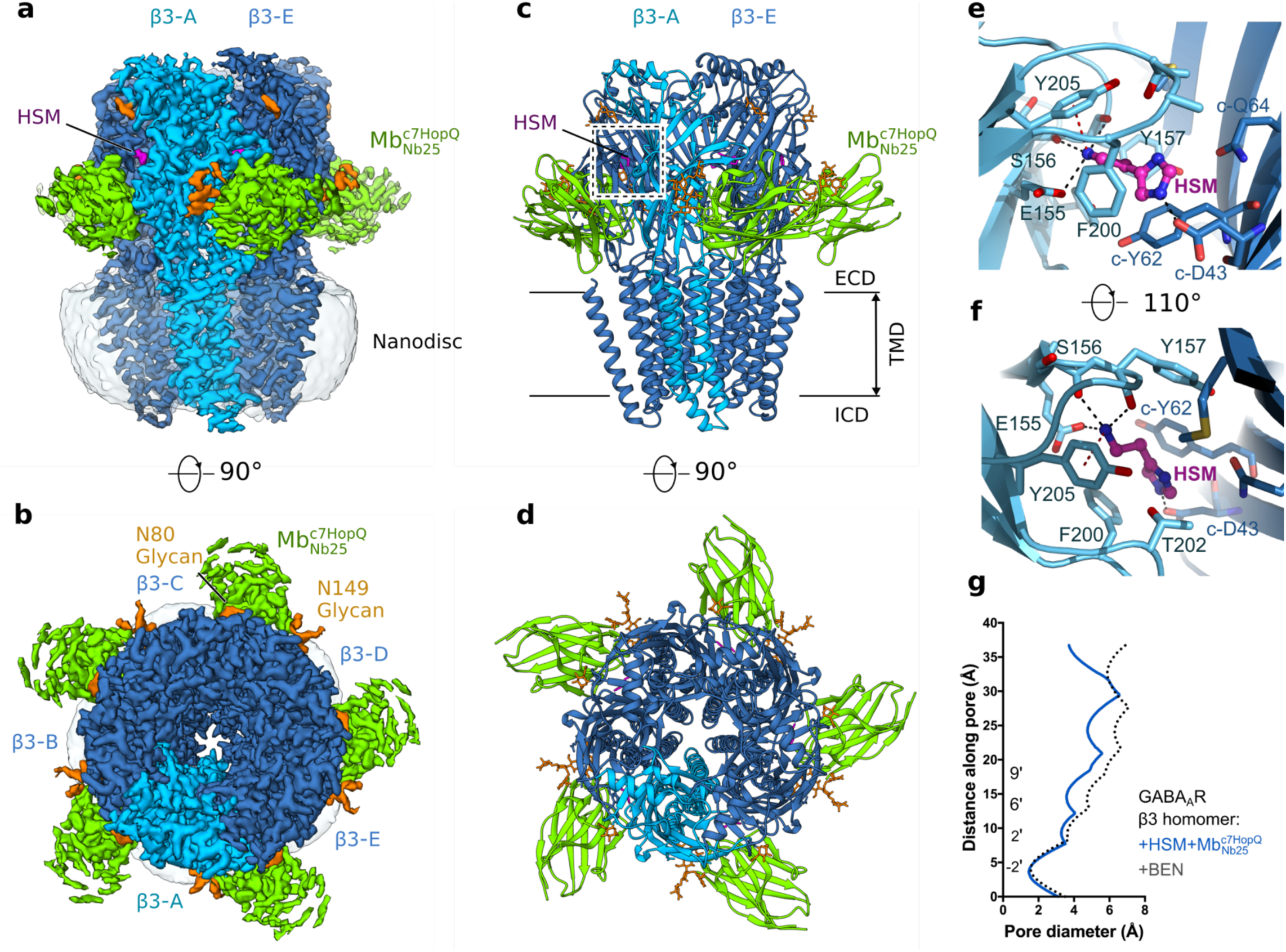
Megabody-enabled high-resolution structure of homopentameric β3 GABA_A_ receptor in lipid nanodiscs. **a-d**, Side (**a**) and (**b**) top view of the sharpened cryo-EM density map of histamine-bound β3 GABA_A_ receptor in a complex with 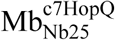 in lipid nanodisc (EMDB-4542, PDB ID: 6QFA). Five β3 subunits are coloured light blue (subunit A) and dark blue (subunits B-E). Histamine (HSM), 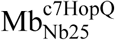 and glycans are coloured magenta, green and orange respectively. Sharpened density maps are contoured at 0.08. Side view (**c**) and top view (**d**) of atomic model of the β3 GABA_A_R in a complex with histamine. **e-f**, Histamine binding mode. Complementary face residues from E subunit marked by ‘c-’. Black and red dashed lines indicate putative hydrogen bonds and cation-π interactions, respectively. **g**, Pore diameter of histamine (HSM) bound β3 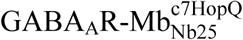 complex (blue line), benzamidine (BEN) bound β3 GABA_A_R (black dotted line, PDB ID: 4COF).

To verify whether the same design principles are not restricted to nanobodies but can also be applied to other antigen-binding domains, we enlarged the small monobody NS1 that binds to both GTP- and GDP-bound states of H-RAS and K-RAS^29^. Monobodies are engineered derivatives of the fibronectin type III domain fold, composed of a seven-stranded β-sandwich built from one three- and one four-stranded β-sheet^25^ (**Supplementary Fig. 7c**). The megabody 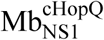 was generated by the insertion of the cHopQ scaffold into the first β-turn (connecting β-strands A and B) of the monobody NS1 (**Supplementary Fig. 7d, e**). Antigen binding was confirmed by measuring binding of fluorescently labelled K-RAS using yeast surface display and flow cytometry^30^ (**Supplementary Fig. 7f**).

### Selection of megabodies from nanobody-immune libraries by yeast display

The complementarity-determining regions (CDRs) which determine the specificity of a given nanobody, are encoded by three hypervariable regions (HV), whereas four more conserved framework (FR) regions define the tertiary structure by encoding nine β-sheets^23,31^ (**Supplementary Fig. 8a**). Building on the structural conservation of the immunoglobulin fold, we found that any given nanobody can easily be reformatted into a megabody of choice by substituting the nanobody sequence, once the connecting linkers between one representative nanobody and a particular scaffold protein are defined. Furthermore, we hypothesised that *in vivo* matured nanobody immune libraries can be cloned as megabody display libraries and screened by conventional phage display or yeast display for the direct selection of functional megabodies. To test this concept, we amplified the open reading frames of the nanobody repertoire of a GFP-immunised llama, lacking the 5’-end that encodes a highly conserved β-strand A. These fragments were cloned in a yeast display vector according to **Fig. 2a** using the same sequence for β-strand A and the same peptide linkers to fuse nanobody to cHopQ as in 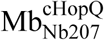 described above (**Fig. 2a, Supplementary Fig. 8**). The generated megabody library was displayed on yeast cells in frame with the Aga2p anchor protein followed by an acyl carrier protein (ACP) for orthogonal labelling^30^ (**Fig. 2b**). This approach enables the identification and subsequent sorting of yeast cells that display high levels of a particular megabody (high DY-647P1 fluorescence) and bind the cognate antigen (high GFP fluorescence) (**Fig. 2c**). After three rounds of selection, nine novel GFP-specific megabodies were identified (**Supplementary Fig. 8**), demonstrating that antigen-binding chimeric proteins can be selected from displayed megabody libraries assembled from llama immune libraries in a straightforward manner.

### Megabodies randomize the orientation of GABA_A_ receptor particles in ice and enable high resolution cryo-EM reconstructions

GABA_A_ receptors (GABA_A_Rs) are pentameric ligand-gated ion channels (pLGICs), which mediate fast inhibitory signalling in the human brain and are targets for clinically-relevant drugs including benzodiazepines and general anaesthetics^32^. In single particle cryo-EM, GABA_A_Rs and related pLGICs are known to suffer from severe preferential orientation in free-standing ice unless detergents are added to the samples. Although such approach in combination with Fab antibody fragments enabled structural analyses of heteromeric GABA_A_ receptors by cryo-EM^33,34^, the presence of detergents damaged the integrity of transmembrane domains (TMDs) and of the interfaces between TMDs and extracellular regions^35^.

To assess the utility of megabodies for single particle cryo-EM and to enable structural studies of human GABA_A_Rs in lipid bilayers, we enlarged nanobody Nb25, which interacts with the extracellular domain (ECD) of the GABA_A_R β3 subunit^36^ to two megabody variants, 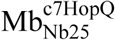 and 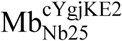, following the same reformatting principals used for Nb207 (**Supplementary Fig. 2a-c**). We inserted a GABA_A_R-β3^37^ homomeric receptor construct into lipid nanodiscs using “on-beads” reconstitution^38,39^ and subsequently vitrified these particles on electron microscopy grids alone or in complex with Nb25, 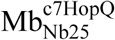 or 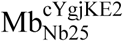. We collected and analysed small cryo-EM datasets for these samples in order to evaluate the extent of preferential particle orientation in each condition (**Fig. 3**). Firstly, when GABA_A_R-β3 homomer samples were frozen alone, ∼84% of the particles observed had their five-fold symmetry axis perpendicular to the water-air interface (“top” views, **Fig. 3a**). Addition of Nb25 did not improve particle orientation (**Fig. 3b**). In both cases, 3D reconstruction attempts led to severely anisotropic maps with large missing regions and low particle distribution efficiencies^15^ (E_od_= 0.18 and 0.22, respectively) (**Fig. 3a,b)**. Remarkably however, more than ∼77% of classified particles displayed different orientations (“tilted” views) when the GABA_A_R-β3 receptor was bound to 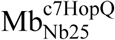 (**Fig. 3c)**. This in turn allowed the reconstruction of a 3D map with a three-fold higher angle distribution efficiency (E_od_= 0.76) and mostly isotropic features. In contrast, 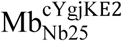 had an opposite effect and induced almost exclusively “top” GABA_A_R-β3 views which led to the most anisotropic 3D reconstruction (**Fig. 3d)**. Megabodies therefore can directly affect the orientation of particles in ice, however it is clear that multiple megabody variants, with different scaffolds, should be tested for a protein of interest.

Based on these findings, we extended this approach to enable cryo-EM studies of the major human heteromeric GABA_A_R variant α1β3γ2. Unlike the homomeric GABA_A_R-β3^37^, heteromeric receptor versions, especially in a physiologically meaningful lipid environment, are intractable for structural analysis by X-ray crystallography. We enlarged the nanobody Nb38 that binds α1 subunits between non-adjacent ECD interfaces^40^ to generate the megabody 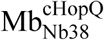, which enabled high-resolution cryoEM structures of the full-length α1β3γ2 human GABA_A_R in lipid nanodiscs bound to a series of small molecule modulators including the competitive antagonist bicuculline, the channel blocker picrotoxin, the agonist GABA and the classical benzodiazepines alprazolam (Xanax) and diazepam (Valium), respectively^38,39^.

### Histamine binding mode and conformational impact on the GABA_A_R-β3 homomer

The 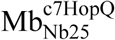 megabody allowed us to solve a 2.49 Å resolution cryo-EM structure of the GABA_A_R-β3 homopentamer bound to its agonist histamine^41^ (**Fig. 4, Supplementary Fig. 9-11, Supplementary Table 3, Supplementary Video 1**). The scaffold region of 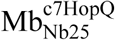 is not visible in the final reconstruction due to the moderate flexibility of its link to the nanobody domain (**Supplementary Fig.12**). However, the high resolution and excellent quality of the cryo-EM map enabled straightforward model building and refinement as well as unambiguous identification of the histamine density, a molecule composed of only eight non-hydrogen atoms. Histamine occupies all five canonical agonist pockets under the sensor loop-C (**Fig. 4c, d, Supplementary Fig. 11d**). The imidazole ring of histamine points towards the complementary (β3^−^) side of the pocket where it is stacked between the side chains of Phe200 and Tyr62 and forms a hydrogen bond with Asp43 side chain (**Fig. 4e, f**). The histamine ethylamine group faces the principal (β3^+^) side of the pocket and forms hydrogen bonds with the Glu155 side chain and backbone carbonyls of Ser156 and Tyr157, and a cation-π interaction with the aromatic ring of Tyr205 (**Fig. 4e, f**). As expected for a GABA_A_R agonist, histamine binding induces closure of the sensor loop-C and an anti-clockwise rotation of each subunit ECD relative to its corresponding TMD, reminiscent to the impact of benzamidine on the GABA_A_R-β3 homopentamer^37^ and GABA on the α1β3γ2L GABA_A_R hetermomeric receptor^39^ (**Supplementary Fig. 10).** The GABA_A_R-β3 channel pore is in a desensitised conformation, with the 9’Leu activation gate open and the −2’ desensitization gate closed, a conformation reminiscent of the benzamidine-bound crystal structure solved in detergent^37^ (**Fig. 4g**). However, unexpectedly, the upper half of the pore is relatively narrower, perhaps reflecting the impact of the lipid environment in which the GABA_A_R-β3 construct described here was reconstituted.

## CONCLUSIONS

We describe here the generic megabody technology, which allows the rapid reformatting, or direct library selection, of high affinity binding domains such as nanobodies or monobodies into larger, conformationally and functionally stable chimeras. We illustrate this technology by using two scaffold proteins, but the same design principle can be applied to any scaffold of any size or geometry required to avoid sterical clashes with the target, providing that it contains a rigid β-hairpin loop that tolerates globular domain insertions. This simple but powerful approach expands the nanobody toolkit for protein structure determination by single-particle cryo-EM but also, at least theoretically, X-ray crystallography because the relatively large and rigid scaffolds themselves offer novel opportunities for crystal contacts. The primary benefit of megabodies in cryo-EM is indeed the ability to randomise the particle distribution in ice for very difficult membrane protein samples, such as pLGICs, that normally exhibit severe preferential orientation. We found cHopQ-based megabodies to be very efficient tools for homomeric and heteromeric GABA_A_ receptors, however the same nanobodies built into a cYgjKE2 scaffold did not help in this case. Therefore, it is possible that the optimal megabody scaffold will vary for other target proteins as well.

Besides the impact on particle orientation we report, the megabodies (∼56 kDa and ∼100 kDa) also facilitate the *in silico* “purification”^42^ of particles, sorting those that contain the stable antigen:megabody complex from those that contain the antigen only or other contaminants. When the nanobody recognition domain is specific to a particular conformation of the target protein, this particle classification step will ensure that the 3D reconstruction obtained represents the antigen in the particular state trapped by the megabody. The increased particle size of antigen:megabody complexes can also facilitate their identification and picking from low-contrast micrographs. Most importantly, the megabody technology has the potential to further stimulate the rapidly growing field of drug discovery using single particle cryo-EM^43^ by enabling high resolution structural analysis of difficult yet highly valuable targets such as eukaryotic membrane proteins.

## ACKNOWLEDGMENTS

We thank A.V. Shkumatov and R.K. Singh for support with SAXS experiments; H. De Greve for providing the GFP+ expressing *E.coli* strain; S. Chen, G. Cannone, G. Sharov and A. Yeates for support at the MRC-LMB EM facility; J. Grimmett, T. Darling and T. Pratt for help with IT and high-performance computing. We thank Instruct-ERIC, part of the European Strategy Forum on Research InfK-ras-rastructures (ESFRI), Instruct-ULTRA (EU H2020 Grant 731005), and the Research Foundation - Flanders (FWO) for support with Nanobody discovery and for funding the PhD training of T.U. Cryo-EM work was supported by the UK Medical Research Council grants MR/L009609/1 and MC_UP_1201/15 to A.R.A.

## AUTHOR CONTRIBUTIONS

E.P. and J.S. conceived the project. T.U. cloned, generated and produced megabodies. A.W. and T.Z. participated in crystallography. T.U. crystallised, H.R. collected and B.F. processed X-ray diffraction data of 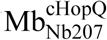. V.K. performed TSA and SAXS experiments. S.M. produced β3 GABA_A_R. S.M. and T.U. collected and processed the electron microscopy data. B.F. and A.W. purified and labelled KRAS. T.U. performed yeast display selection. W.V. implemented the computational modelling. T.U., A.R.A. and J.S. wrote the manuscript. All authors participated in discussion and revision of the manuscript.

## METHODS

### Construction, expression and purification of 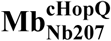

pMESD2 (**Supplementary Fig. 2**), a versatile plasmid for the cloning and the expression of megabodies that are built from nanobodies that are grafted into a circularly permutated variant of the extracellular adhesin domain of *H. pylori* (HopQ, UniProtKB B5Z8H1), is a derivative of pMESy4 (GenBank KF415192). The open reading frame of pMESD2 is under the transcriptional control of the *Plac* promotor and encodes the DsbA leader sequence, followed by a consensus sequence encoding the conserved thirteen residues of β-strand A of the nanobody fold^44^ (residues 1-14 in IMGT^45^ numbering, QVQLVESGGGLVQ), followed in frame by the C-terminal part of HopQ (residues 227-449, UniProtKB B5Z8H1), followed by a short peptide linker ASGGGSGGGGSG connecting the C-terminus and the N-terminus of HopQ to produce a circular permutant of the scaffold protein, followed by the N-terminal part of HopQ (residues 49-221, UniProtKB B5Z8H1), followed by a Gly residue (residue 17 of a Nanobody), followed by a multi cloning site (MCS), followed by the His6 tag and the EPEA tag^46^. To clone and express 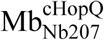, a gene fragment encoding β-strands B to G of a GFP-binding nanobody Nb207 (residues 18-128 in IMGT numbering) was amplified by PCR using primers TU89 and EP230 and cloned as a SapI fragment in pMESD2, enabling the expression and secretion of the 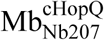 to the periplasm of *E. coli*. To express 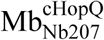, WK6 cells^47^ bearing the pMESD2_ 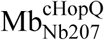 plasmid were grown in Terrific Broth medium supplemented with ampicillin (100 mg/ml) at 120 rpm and 37 °C to OD_600_ = 4, and induced overnight with 1mM IPTG at 28 °C. Cells were harvested by centrifugation (5,000*g*, 15 min) and the recombinant 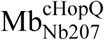 was released from the periplasm using an osmotic shock^24^. The megabody was next separated from the protoplasts by centrifugation and recovered from the clarified supernatant on a HisTrap FF 5mL prepacked column. The protein was next eluted from the Ni-NTA resin by applying 500 mM imidazole and concentrated by centrifugation using Amicon Ultra Filters (cut-off of 10 kDa, Sigma). Concentrated samples were applied on a Superdex 200 PG 16/90 size exclusion column (GE Healthcare) equilibrated with 10 mM Tris pH 7.3 and 140 mM NaCl and eluted as a single peak (**Supplementary Fig. 2**).

### Structure determination of 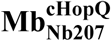 by X-ray crystallography

Purified 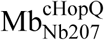 was concentrated to 48 mg/mL and subjected to a number of commercial sparse-matrix crystallisation screens (JSCG/Proplex/PEGion/Wizard12/Morpheus) in 0.1 µL sitting drops supplemented with 0.1 µL of the mother liquor. Small crystals were obtained in the A2 condition (0.1 M sodium citrate, pH 5.5, 20% w/v PEG3000) of the JSCG screen. Well-diffracting crystals were grown by seeding 0.2 M ammonium citrate, 17% PEG3350, 10% glycerol and 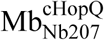 at 48 mg/mL with the JSCG A2 crystals. Data were collected at 100K on the I03 source at Diamond Light Source synchrotron (Oxfordshire, UK) and the structure was refined to 2.84 Å resolution. The Megabody crystallised in P1 with 10 molecules in the asymmetric unit. Diffraction data were integrated and scaled with XDS^48^. Models were built by iterative cycles of refinement with Phenix and Buster-TNT^49^ and manual building in Coot^50^. MolProbity was used for structure validation^51^. Data collection and refinement statistics are summarised in **Supplementary Table 2**. Root mean square deviations (RMSD) and rotation angles were calculated using UCSF Chimera^52^.

### Small-angle X-ray scattering of 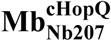

We collected small-angle X-ray scattering (SAXS) data for 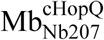 in a mail-in session at the B21 beamline of Diamond Light Source synchrotron (Oxfordshire, UK). The scattering intensities were recorded in a Size-Exclusion Chromatography coupled SAXS (SEC-SAXS) experiment after injection of the protein sample on a Superdex 200 Increase 3.2/300 gel filtration column (GE Healthcare) equilibrated with 10 mM Tris-HCl pH 7.3, 14 mM NaCl on an Agilent HPLC instrument. The averaging and buffer subtraction of the resulting data frames were performed using the program DATASW^53^. The data corresponding to the peak of the protein were further processed using the ATSAS software package^54^. The scattering curve was generated with PRIMUS and was subjected to indirect Fourier transform using GNOM to yield the pair-distance distribution function P(r), from which the radius of gyration (*R*_*g*_) and the maximum particle dimension (*D*_*max*_) were estimated. R_g_ was obtained also from the slope of the Guinier plot in PRIMUS. Further interpretation of the SAXS data involved using the dimensionless Kratky plot. The theoretical scattering profile of the X-ray structure of 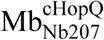 was calculated using CRYSOL^55^.

### Construction, expression and purification of 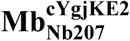

pMESP3E2 (**Supplementary Fig. 2**), a versatile plasmid for the cloning and the expression of megabodies that are built from nanobodies grafted into a circularly permutated variant of the *E.coli* K12 glucosidase YgjK (UniProtKB P42592), is a derivative of pMESy4 (GenBank KF415192). The open reading frame of pMESP3E2 is under the transcriptional control of the *Plac* promotor and encodes the PelB leader sequence^56^, followed by a consensus sequence encoding the conserved twelve residues of β-strand A of the Nanobody fold^44^ (residues 1-13 in IMGT numbering, QVQLVESGGGLV), followed in frame by one additional Tyr residue, followed by the C-terminal part of YgjK (residues 487-783, UniProtKB P42592), followed by a short peptide linker ASGGGSGGGGSGGGGSG connecting the C-terminus and the N-terminus of YgjK to produce a circular permutant of the scaffold protein, followed by the N-terminal part of YgjK (residues 24-484, UniProtKB P42592), followed by one additional Asp residue, followed by a multi cloning site (MCS), followed by the His6 tag and the EPEA tag^57^. For the expression and secretion of 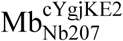 to the periplasm of *E. coli*, the gene fragment encoding β-strands B to G of Nb207 (residues 18-128 in IMGT numbering) was amplified by PCR using primers TU89 and EP230 and cloned as a SapI fragment in pMESP3E2. WK6 cells^47^ bearing the 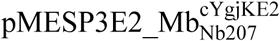 plasmid were inoculated in Terrific Broth medium supplemented with ampicillin (100 mg/ml), grown at 120 rpm and 37 °C to OD_600_= 4 and induced overnight with 1mM IPTG at 28 °C. Cells were harvested by centrifugation (5,000*g*, 15 min) and 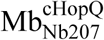 was released from the periplasm by resuspending the cells in 20% w/v sucrose supplemented with 0.5 mg/ml lysozyme (L6876 Sigma), 50 mM Tris pH 8, 1 mM EDTA and 150 mM NaCl for 30 min at 4 °C followed by centrifugation for 30 min at 10,000*g* to separate the protoplasts. The supernatant was supplemented with 500 mM NaCl and 5 mM MgCl_2_ and applied on a HisTrap FF 5mL prepacked column (GE Healthcare). Proteins were eluted from the resin with 500 mM imidazole and concentrated by centrifugation using Amicon Ultra Filters (cut-off of 50 kDa, Sigma). For polishing, concentrated samples were applied on a Superdex 200 PG 16/90 size exclusion column equilibrated with 20 mM Tris at pH 7.3, 140 mM NaCl buffer and eluted in a single symmetric peak (**Supplementary Fig. 2**).

### Thermal stabilities of Nb207, 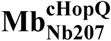 and 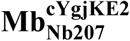

We compared the thermal stabilities of Nb207, 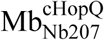 and 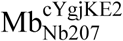 by measuring the increase in fluorescent intensity of the partition hydrophobic-binding dye SYPRO® Orange in the presence of the different proteins upon thermal melting. Measured samples contained 5x SYPRO® Orange in 10 mM Tris pH 7.3, 140 mM NaCl with final protein concentrations 0.2 mg/ml for the nanobody and 2 mg/ml for the megabodies, respectively. Samples for the thermal shift assays were prepared in triplicate (n=3) in a 96-well qPCR microplate (BioRad) in a final volume of 20 µL. Thermally-induced protein melting was performed in a CFX qPCR instrument (BioRad) using a temperature gradient from 25 to 100 °C at a heating rate of 1 °C per minute (**Supplementary Fig. 2**). Experimental data were fitted with GraphPad Prism 7 using Boltzmann’s equation Y=Bottom+(Top-Bottom)/(1+exp((V50-X)/Slope)).

### Expression and purification of GFP

GFP variant GFP+^58^ was expressed in *E.coli* strain DH5 alpha^59^ as a C-terminally His6-tagged protein under the transcriptional control of the *lac* promoter using pUC8. Cells were grown overnight in Luria-Bertani broth supplemented with ampicillin (100 mg/ml) at 37 °C, harvested by centrifugation (5,000*g*, 15 min), resuspended in lysis buffer (50 mM Tris pH 8, 200 mM NaCl, 15 mM EDTA) and lysed with a high-pressure homogeniser (Constant System). Lysed cells were next pelleted by centrifugation for 30 min at 10,000*g* and the supernatant was applied on a HisTrap FF 5mL prepacked column (GE Healthcare). GFP was eluted with 500 mM imidazole and concentrated by centrifugation using Amicon Ultra Filters (cut-off of 3 kDa, Sigma) and polished on a Superdex 75 PG 16/90 size-exclusion column equilibrated with 20 mM Tris pH 7.3 and 140 mM NaCl.

### Expression, purification and fluorescent labelling of KRASG12V

A codon-optimised synthetic gene encoding the human KRASG12V mutant (residues 1-169) was cloned as an NdeI and XhoI fragment into pET28b (Novagen). His-tagged K-RASG12V, abbreviated KRAS in this paper, was expressed in *E. coli* strain BL21 and purified as described previously^60^. For applications in FACS, purified KRAS was labelled with a five-fold molar excess of the DyLight650-NHS ester (Thermo Fisher Scientific) following the manufacturer’s instructions. After a 30 min incubation at room temperature, unreacted label was quenched with 50 mM Tris pH 8.0 and the labelled protein was separated from free label by size-exclusion chromatography on a Superdex 200 PG 16/90 column (GE Healthcare).

### Binding kinetics of Nb207, 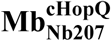 and 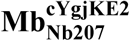 to GFP

We used bio-layer interferometry (BLI) on an OctetRED96 (ForteBio) to compare the binding kinetics of Nb207, 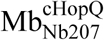 and 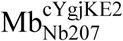 to GFP. For immobilisation on the biosensors, purified GFP was biotinylated with a five-fold molar excess of EZ-link NHS-Biotin (Thermo Fisher Scientific) following the manufacturer’s instructions and separated from unreacted NHS-biotin on a NAP10 column (GE Healthcare). The biotin/GFP ratio was determined at 2.08 ± 0.02 using the Pierce Biotin Quantitation kit (Thermo Fisher Scientific). For BLI, biotinylated GFP was diluted to 0.75 µg/ml in 10 mM Tris pH 7.3, 140 mM NaCl and 0.05% Tween20 supplemented with 1 mg/mL BSA and directly immobilised on Streptavidin (SA) biosensors at about 1 nm response. After two equilibration steps of 100 s, the binding isotherms were monitored by exposing separate sensors simultaneously to different concentrations of Nb207, 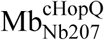 or 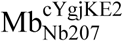, respectively. Association kinetics were followed for 300 s at 30 °C under constant stirring at 1000 rpm, tailed by dissociation experiments for 2800 s. Association and dissociation rates were estimated by fitting the sensograms using the 1:1 binding model included in the Octet Data Analysis software 9.1 (ForteBio).

### Selection of GFP-specific megabodies from Nanobody-immune libraries by yeast display and FACS

Plasmid pNMB2 is a derivative of yeast surface display vector pNACP^30^. A new display cassette encoding the following elements was synthesised and cloned as an EcoRI-BglII fragment to replace the original cassette: appS4 leader sequence (LS) for secretion^61^, followed by a consensus sequence encoding the conserved β-strand A of the nanobody fold^44^ (residues 1-14 in IMGT numbering, QVQLVESGGGLVQ), the C-terminal part of HopQ^26^ (residues 227-449, UniProtKB B5Z8H1), a short peptide linker ASGGGSGGGGSG connecting the C-terminus and the N-terminus of HopQ to produce a circular permutant of the scaffold protein, the N-terminal part of HopQ (residues 49-221, UniProtKB B5Z8H1), a multi cloning site (MCS), the Aga2p anchor protein followed by the Acyl Carrier Protein (ACP) and the Myc tag (**Fig. 2a**). The parental cloning vector pNMB2 encodes an in frame stop codon in the MCS to avoid background display and orthogonal staining of ACP from plasmids that do not contain an insert.

A llama was immunised with GFP+^58^ and mRNA fragments encoding the full collection of the naturally circulating nanobody repertoire of the immunised animal were isolated as described^24^. Briefly, total RNA was isolated from the peripheral blood lymphocytes (PBLs) of the immunised animal to prepare cDNA and the open reading frames encoding all immunoglobulin heavy-chains were amplified by RT-PCR with primers call001 and call002. Fragments encoding the nanobodies from β-strands B to G were amplified thereof through a nested PCR using primers TU64 and TU65 (**Fig. 2a, Supplementary Fig. 8, Supplementary Table 4**). 10 µg of this PCR product was mixed with 10 µg of BamHI/HindIII linearised pNMB2 and transformed into electrocompetent EBY100 *Saccharomyces cerevisiae* cells for GAP repair homologous recombination^62^ to produce a library of yeast cells that display diverse cHopQ megabodies derived from the nanobody repertoire of a GFP-immunised llama.

This yeast library displaying diverse megabodies was inoculated, induced and orthogonally stained with CoA-647 to monitor the display level of the Mb-Aga2p-ACP fusion on each yeast cell as described^30^. In each round of selection, 4 × 10^7^ of these stained yeast cells were incubated with GFP for 60 min at 4 °C in 500 µl of cold PBS supplemented with 0.2% (w/v) BSA at pH 7.4 (PBS–BSA). Yeast cells were next washed three times with and resuspended in 2 ml of PBS–BSA and sorted on a FACS Aria (BD Biosciences). Selected yeast cells were recovered into SDCAA medium, grown and induced, then stained again for subsequent rounds of selection. Individual yeast clones expressing a GFP binding megabody were grown separately in 96-well plates for sequencing and further characterisation as described^30^. Briefly, each clone was confirmed to become green fluorescent after incubation with 100 nM GFP followed by 3 wash steps by flow cytometry using FACS Fortessa (BD Biosciences). Routinely, ∼10,000 yeast cells derived from a particular clone were analysed to calculate the GFP mean fluorescence intensity (MFI) using FlowJo software (FlowJo, LLC) and compared to the fluorescence of a yeast clone displaying an irrelevant megabody as the negative control.

### Expression and purification of Nb25, 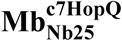 and 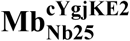

Nb25 was expressed and purified as described before^36^. pMESD2c7 is a variant of cloning vector pMESD2 in which the C-terminal end of HopQ scaffold protein (residues 227-446, UniProtKB B5Z8H1) is directly fused to its N-terminal end (residues 53-221, UniProtKB B5Z8H1) omitting the additional flexible linker to implement circular permutation. We constructed this alternative circular permutation variant of HopQ (c7HopQ) because the linker used to design our first megabody 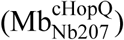 was flexible and not visible in the electron density maps (PDB ID: 6QD6). The gene fragment encoding β-strands B to G of the β3 GABA_A_R-specific Nb25^36^ (residues 18-128 in IMGT numbering) was amplified by PCR using primers TU89/EP230 (**Supplementary Table 4**) and cloned as a SapI fragment into pMESD2c7 or pMESP3E2 to produce 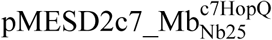 and 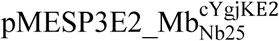, respectively. All Nb25-derived megabodies were expressed and purified as described above.

### Expression of human β3 homopentameric GABA_A_R

A human β3 homopentameric GABA_A_R (UniProtKB P28472) construct which contains the K279T point mutation for increased stability, an SQPARAA linker^32^ substituting the M3-M4 loop, and a C-terminal (GGS)_3_GK–Rhodopsin-1D4-tag (TETSQVAPA)^63^ was transiently expressed in HEK293S-GnTI^-^ cells as described^37^. Briefly, HEK293S-GnTI^-^ cells were grown in protein expression medium (PEM, Thermo Fisher Scientific) supplemented with 1% foetal bovine serum (Invitrogen) at 37 °C, 8% CO_2_. Cells were transfected with the DNA-PEI mix at a ∼2×10^6^ cells/ml density, and 48 h post-transfection were harvested by centrifugation at 4,000*g*, 4 °C. Cell pellets were snap-frozen in liquid N2 and stored at −80 °C for future use.

### Nanodisc reconstitution of human β3 homopentameric GABA_A_R

All purification and reconstitution steps were performed at 4 °C or on ice. Each of three cell pellets from 0.8 l culture were resuspended by vortexing in dilution buffer: 50 mM HEPES pH 7.6, 300 mM NaCl, 1 mM histamine, 1% (w/v) mammalian protease inhibitor cocktail (Sigma-Aldrich). Solubilisation was performed for 1 h by adding 1% (w/v) lauryl maltose neopentyl glycol (LMNG, Anatrace) and cholesterol hemisuccinate (CHS, Anatrace) at a 10:1 (w/w) ratio. Solubilised GABA_A_R was separated from insoluble material by centrifugation (10,000*g*, 15 min) and captured on a 1D4 affinity resin (250 μl) by slow rotation for 2 h. The resin was harvested (300*g*, 5 min) and washed three times with 50 ml of washing buffer: 50 mM HEPES pH 7.6, 300 mM NaCl, 1 mM histamine (Sigma-Aldrich), 1% (w/v) LMNG and 0.1% CHS. The washed resin was equilibrated with 1 ml of dilution buffer and 240 μl of a mixture containing 80% (w/v) phosphatidylcholine (POPC, Avanti) and 20% of a bovine brain lipid (BBL) extract (Sigma-Aldrich). After 30 min incubation, the resin was equally divided in five Eppendorf tubes and collected by centrifugation. For nanodisc reconstitution, Bio-Beads (10 mg/ml final concentration) with an excess of MSP2N2^64^ (0.6 mg ml^-1^ final concentration) were added to each sample. 100 μl of Nb25, 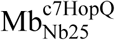 and 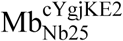 (∼120 μM) were added to corresponding sample tubes and incubated for 1 h slowly rotating. Resin samples were harvested (300*g*, 5 min), washed six times with dilution buffer, resuspended in 50 μl of elution buffer: 12.5 mM HEPES pH 7.6, 75 mM NaCl, 0.25 mM histamine, 1.5 mM 1D4 peptide (Cube Biotech) and incubated overnight. Beads were pelleted by centrifugation (300*g*, 5 min) to collect the supernatants. These supernatants were supplemented once more with 0.4 μl of ∼ 120 μM of Nb25, 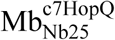, and 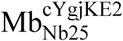, respectively to be used for cryo-EM grid preparation.

### Cryo-EM sample preparation, image collection and processing

We used the same batch of β3_K279T_ GABA_A_R reconstituted in lipid nanodiscs to prepare cryo-EM grids of the receptor alone, in complex with Nb25, 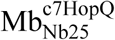 or 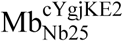, respectively. 3.5 μl of each sample was applied onto glow-discharged gold R1.2/1.3 300 mesh UltraAuFoil grids (Quantifoil) for 30 s and blotted for 5.5 s before vitrification in liquid ethane. A Vitrobot Mark IV (Thermo Fisher Scientific) was used for plunge-freezing at ∼100% humidity and 14.5 °C.

Cryo-EM data of all samples were collected on a 300 kV Titan Krios microscope (Thermo Fisher Scientific) using a Falcon 3EC (Thermo Fisher Scientific) direct electron detector in counting mode and a Volta Phase Plate (VPP, Thermo Fisher Scientific). Data collection parameters are provided in **Supplementary Table 3**.

In order to investigate the proportion of preferential particle views of β3 GABA_A_R particles in samples where β3 homomer was alone or complexed with Nb25, 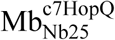 or 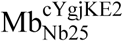, small cryo-EM datasets were analysed by using the same basic data processing procedure. First, MotionCor2^65^ was used to motion-correct the movies and Warp^66^ was applied to estimate the contrast transfer function (CTF), phase shift parameters and to pick, and extract particles. The reference-free 2D classification was performed using RELION 3.0^67^. One round of 2D classification was performed and well-aligned 2D classes showing clear GABA_A_R particle projections were used to determine the proportion of preferred particle orientations in each sample (around 6,000 particles for each of the conditions). Next, the particles from the 2D classification were subjected to reference-free 3D model generation and 3D refinement using cryoSPARC^68^. The efficiency of the particle orientation distribution (E_od_ values) for each 3D model was calculated using cryoEF^15^.

For the high-resolution reconstruction of β3 GABA_A_R bound to 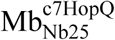, a larger dataset was processed using RELION 3.0 as described below. MotionCor2 and Gctf^69^ wrappers inside RELION 3.0 package were used to motion-correct movies and to estimate the contrast transfer function and phase shift parameters, respectively. Manual inspection was used to discard poor quality movies. Next, particles were auto-picked using a Gaussian blob autopicker function in RELION 3.0. The resulting particles were 2D classified and the best classes were selected for further processing. Stochastic gradient descent (SGD) methodology^68^ (RELION 3.0) was used to generate an initial reference-free 3D model. A ‘gold standard’ 3D-refinement was performed and subjected to Bayesian particle polishing^70^. Next, particles were sorted using 3D classification jobs without particle alignment and the highest-resolution classes were used for a 3D-refinement using a soft mask and solvent-flattened Fourier shell correlations (FSCs). Further beam tilt correction and per-particle contrast transfer function refinement implementations in RELION 3.0 were applied, yielding the final cryoEM map of 2.49 Å resolution (FSC criteria of 0.143, **Supplementary Figure 9**). Local map resolution was estimated with ResMap^71^.

For atomic model generation, the coordinates of β3 GABA_A_R (PDB ID: 4COF) and α5β3 chimera (PDB ID: 5O8F) bound to Nb25 were used as templates. First, the atomic coordinates of β3 homomer ECD, TMD and Nb25 were fitted to the highest resolution map (2.49 Å) as rigid bodies using UCSF Chimera. Then, the coordinates were manually adjusted using COOT^50^ followed by several rounds of global refinement and minimisation in real space with *phenix_real_space_refine*^72^. The geometry constraint files for histamine molecule were generated using Grade Web Server (Global Phasing). The model geometry quality assessment was performed using the MolProbity^51^ web server. To validate the refinement protocol, the coordinates of the final model were displaced by 0.5 Å using CCP-EM software^73^. The resulting model was refined with *phenix_real_space_refine* against one of the half-maps produced by RELION 3.0. FSC curves were calculated between this model and the half-map used for refinement (‘work’) and the half-map, which was not used for refinement, (‘free’) using *phenix.mtriage*^74^. In addition, the FSC curve was calculated for the refined model vs the final sharpened map (‘full’). The separation between FSC_work_ and FSC_free_ curves was not significant, indicating the model was not over-refined. Pore diameters were calculated using the HOLE^75^ plug-in available in Coot.

### Construction and display of 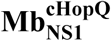 on a yeast

pNS1MB (**Supplementary Fig. 5**) is a derivative of yeast surface display vector pNACP^30^ encoding the following elements: appS4 leader sequence (LS)^61^, β-strand A of monobody NS1^29^ (residues 4-16), an additional Phe residue, C-terminal part of HopQ (residues 227-449, UniProtKB B5Z8H1), a short peptide linker ASGGGSGGGGSG connecting the C-terminus and the N-terminus of HopQ to produce a circular permutant of the scaffold protein, an N-terminal part of HopQ (residues 49-221, UniProtKB B5Z8H1), an additional Gly residue, C-terminal part of monobody NS1 (residues 19-97), the Aga2p anchor protein, the ACP and the Myc tag (**Supplementary Fig. 7**).

Yeast cells containing pNS1MB vector were inoculated and orthogonally stained with CoA-488 to monitor the display level of the 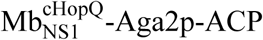 fusions as described^30^. Yeast cells containing 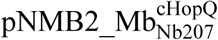 to display 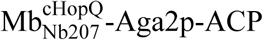 were used in parallel as a control. 10^5^ yeast cells stained with CoA-488 were incubated for 60 min at 4 °C (rotating at 50 rpm) with the KRAS-Dylight650 in 100 µl of cold PBS–BSA. Cells were next washed three times with PBS–BSA, resuspended in 100 µl and applied on FACS Fortessa (BD Biosciences). For ∼10,000 yeast cells displaying 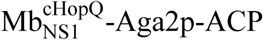 fusion, Dylight650 and ATTO-488 fluorescence was analysed using FlowJo software (FlowJo, LLC) and compared to the negative control.

