## Supplementary Data for "Megabodies expand the nanobody toolkit for protein structure determination by single-particle cryo-EM"

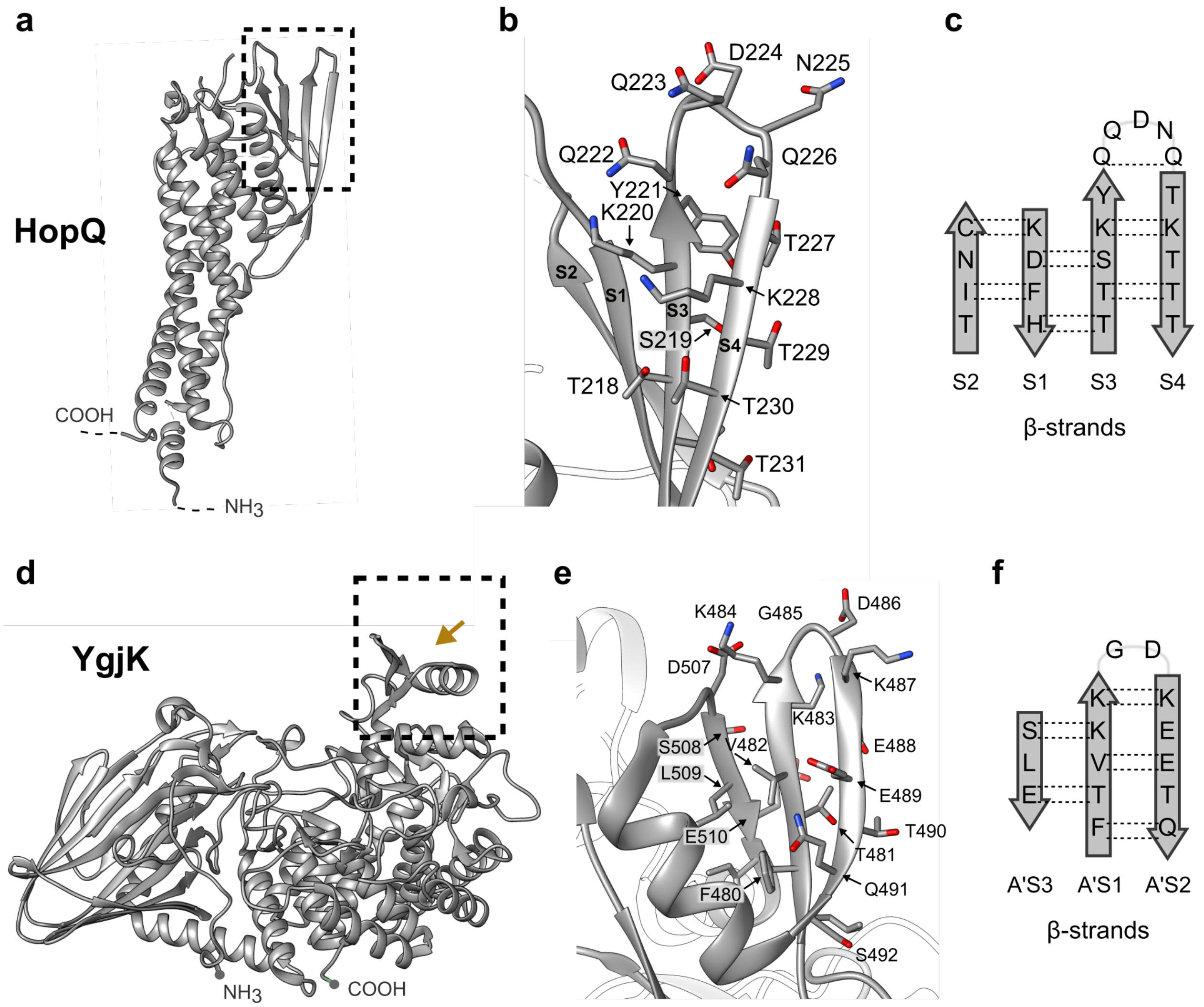

**Supplementary Figure 1. Structures of the scaffold proteins HopQ and YgjK. a-c,** Cartoon representation of the extracellular adhesin domain of *H. pylori* crystal structure (HopQ, PDB ID: 5LP2). **a**, The flexible N- and C-terminal regions are invisible in the electron density and are indicated by dashed lines. The boxed region is enlarged in (**b**). Residues are numbered according to UniProtKB B5Z8H1. **c**, Secondary structure of the solvent-exposed S3-S4 β-turn. Hydrogen bonds between the backbone atoms are indicated by dotted lines. **d-f,** Cartoon representation of the *E. coli* K12 Glucosidase crystal structure (YgjK, PDB ID: 3W7T). **d**, N- and C-termini are indicated by dots. The boxed region is enlarged in (**e**). Residues are numbered according to UniProtKB P42592. **f**, Secondary structure of the solvent-exposed A'S1-A'S2 β-turn.

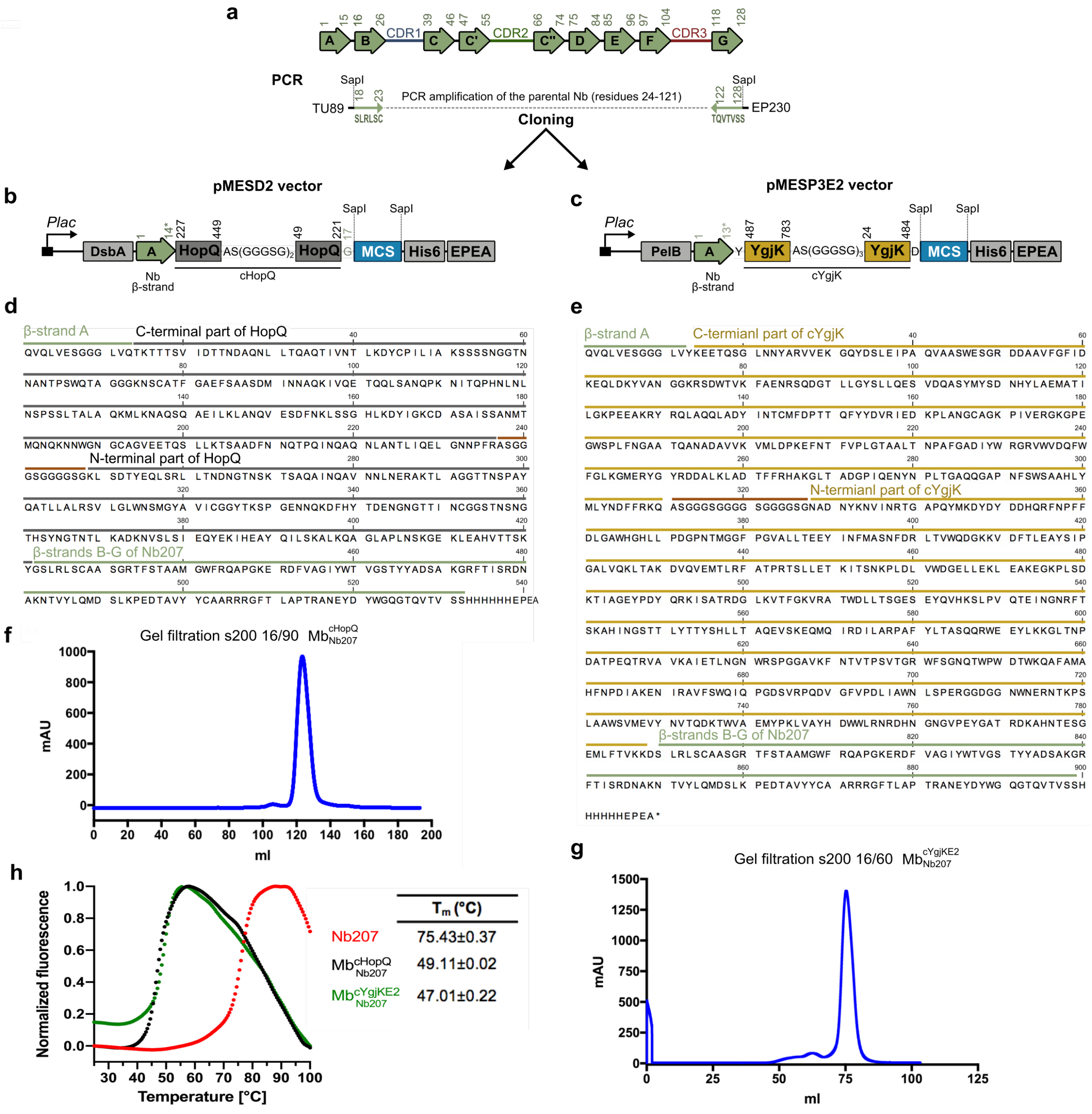

**Supplementary Figure 2. Cloning, expression and purification of megabodies $\mathrm{Mb}^{\mathrm{cHopQ}}_{\mathrm{Nb207}}$ and $\mathrm{Mb}^{\mathrm{cYgjKE2}}_{\mathrm{Nb207}}$. a,** Gene fragments encoding β-strands B to G of a nanobody are amplified by PCR using TU89 and EP230 primers and cloned into pMESD2 (**b**) to turn a nanobody into the cHopQ-megabody format or cloned in pMESP3E2 (**c**) for the cYgjK format. The residues of nanobody, HopQ and YgjK are numbered according to IMGT, UniProtKB B5Z8H1 and UniProtKB P42592. d-e, Amino acid sequences of $\mathrm{Mb}^{\mathrm{cHopQ}}_{\mathrm{Nb207}}$ (**d**) and $\mathrm{Mb}^{\mathrm{cYgjKE2}}_{\mathrm{Nb207}}$ (**e**). **f-g,** Size exclusion profiles (Superdex 200 PG 16/90) of $\mathrm{Mb}^{\mathrm{cHopQ}}_{\mathrm{Nb207}}$ (**f**) and $\mathrm{Mb}^{\mathrm{cYgjKE2}}_{\mathrm{Nb207}}$ (**g**), purified from the periplasm of *E. coli* by Ni-NTA affinity chromatography. **h,** Representative melting curves of Nb207, $\mathrm{Mb}^{\mathrm{cHopQ}}_{\mathrm{Nb207}}$ and $\mathrm{Mb}^{\mathrm{cYgjK}}_{\mathrm{Nb207}}$ measured by thermal shift assays using the partition hydrophobic-binding dye SYPRO® Orange. Experiments were performed in triplicates and the raw data were fitted to the Boltzmann's equation using Prism 7 software (GraphPad) to calculate melting temperatures ($T_m$).

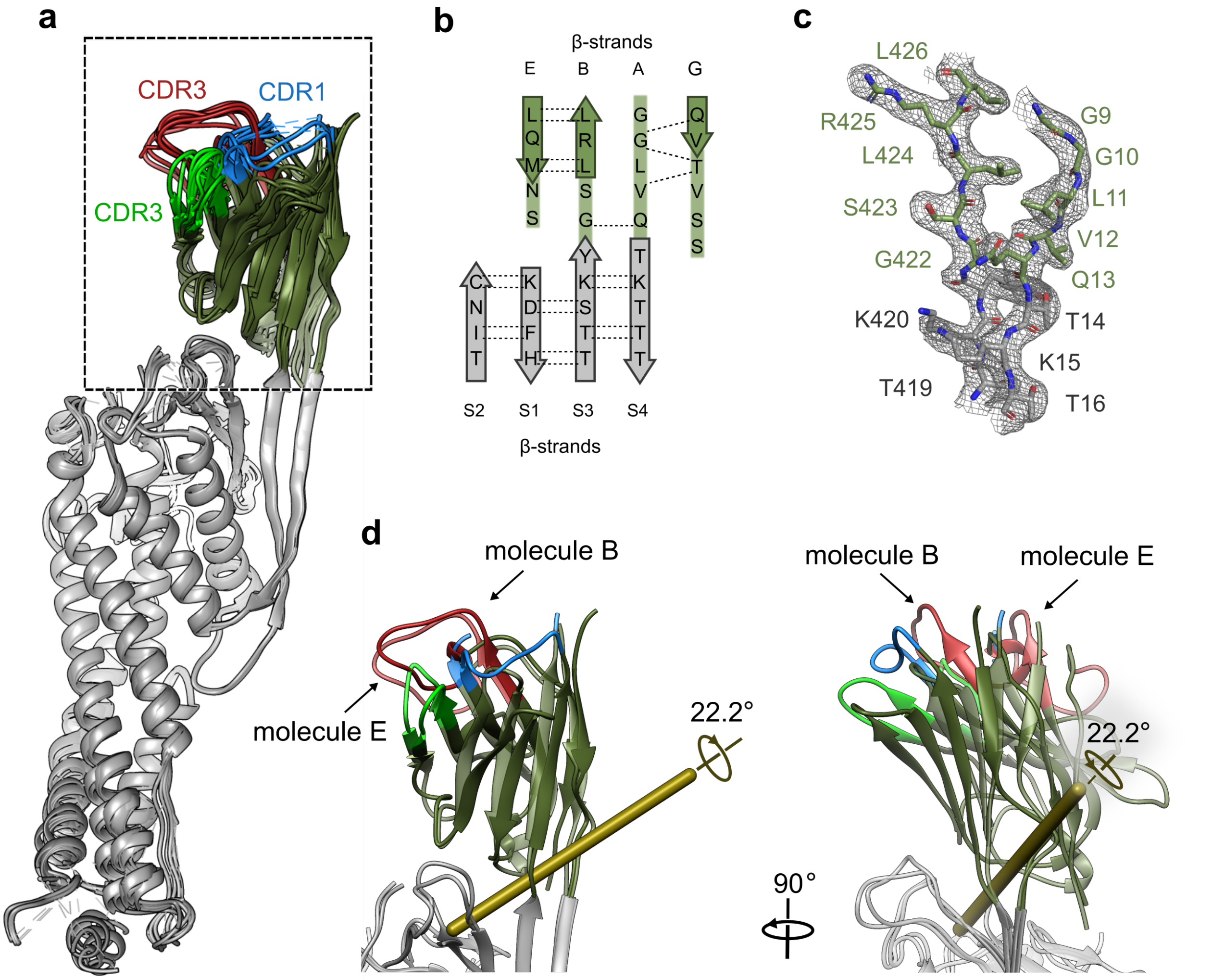

**Supplementary Figure 3. Crystal structure of megabody $\mathbf{Mb^{cHopQ}_{Nb207}}$. a,** Comparison of the ten $Mb^{cHopQ}_{Nb207}$ molecules present in the asymmetric unit (PDB ID: 6QD6). Molecules were aligned using the Cα atoms of the scaffold protein (cHopQ, grey) manifesting minor bending of the Nb207 part (green). **b,** Schematic representation of the β-sheet topology within the region connecting Nb207 to the scaffold. Hydrogen bonds between backbone atoms are indicated by dotted lines. **c,** 2Fo–Fc electron density map (contoured at 1.0 σ) containing the peptides connecting Nb207 (green) to cHopQ (grey) in molecule F. **d**, Structural comparison of molecules B and E that possess the most distinct bending of the nanobody part (residues 1-13 and 422-532 of $Mb^{cHopQ}_{Nb207}$). The rotation axis (gold stick) and angle are calculated for the nanobody part.

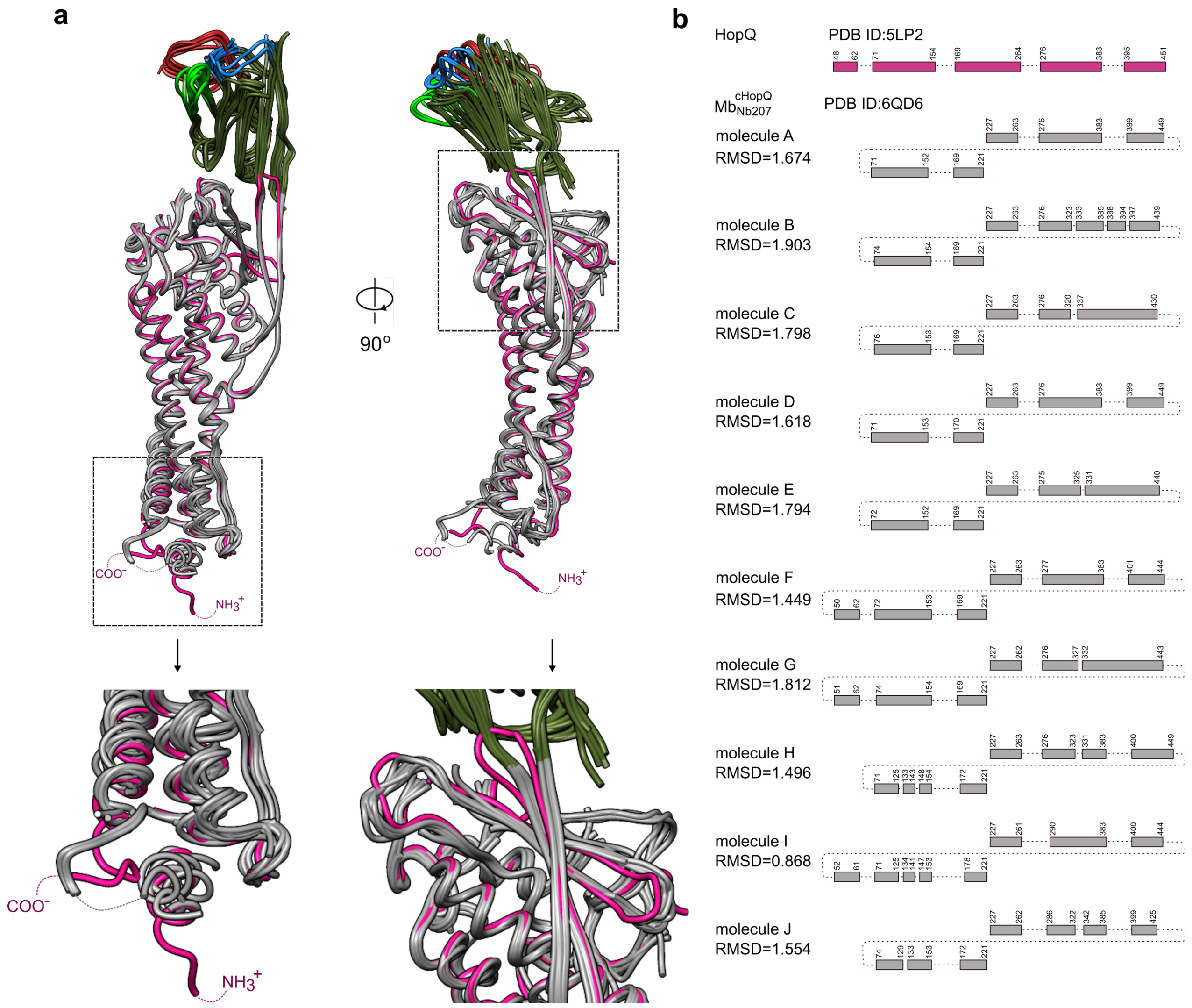

**Supplementary Figure 4. Structural comparison of the parental *H. pylori* adhesion domain to the circularly permutated scaffold in $\mathbf{Mb^{cHopQ}_{Nb207}}$.** **a**, Alignment of each molecule of $Mb^{cHopQ}_{Nb207}$ in the asymmetric unit (coloured in grey-green, PDB ID: 6QD6) onto the *H. pylori* adhesin domain (coloured in magenta, PDB code:5LP2). **b,** The RMSD values between the *H. pylori* adhesin domain (magenta) and the different $Mb^{cHopQ}_{Nb207}$ molecules in the asymmetric unit (grey) were calculated from all corresponding $C_\alpha$ atoms that are refined in the respective electron density maps. The block diagrams describe the segments that are visible/invisible in the adhesin crystal structure and ten megabody molecules in the asymmetric unit, taking into account the circularly permutated arrangement of the scaffold protein.

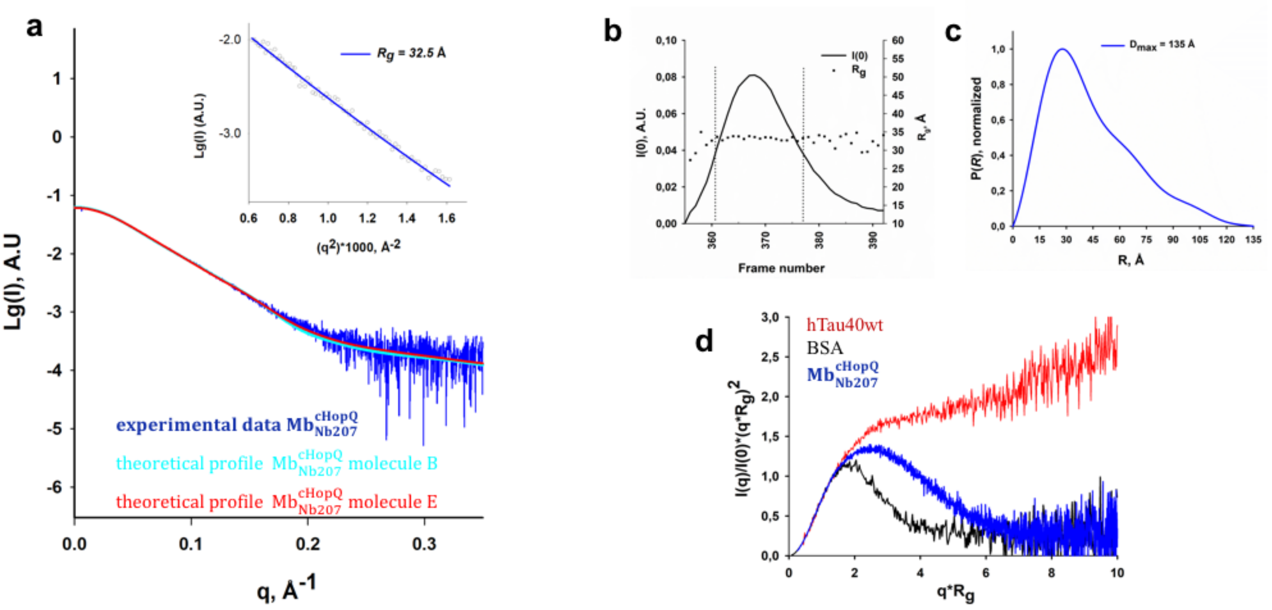

**Supplementary Figure 5. SEC-SAXS analysis of megabody $Mb^{cHopQ}_{Nb207}$. a,** Superposition of the experimental scattering profile of $Mb^{cHopQ}_{Nb207}$ (blue) on the theoretical profiles of molecule B (cyan) and molecule E (red) calculated from the X-ray structure (PDB ID: 6QD6) using CRYSOL ($\chi^2$ = 1.699 for molecule B and $\chi^2$ = 2.033 for molecule E). The inset figure shows the linear Guinier region from the experimental scattering curve and is indicative of a non-aggregated protein sample. The respective $R_g$ value is given. **b,** Elution profile of a SEC-SAXS experiment (black line, frame range of the peak 361-377). A stable $R_g$ is observed over the entire elution profile (black squares). **c,** Normalised P(r) profile with derived $D_{max}$ value. **d,** Dimensionless Kratky plot for $Mb^{cHopQ}_{Nb207}$ (blue) in comparison with two reference proteins: the highly flexible hTau40wt (red) and the globular BSA (black).

a

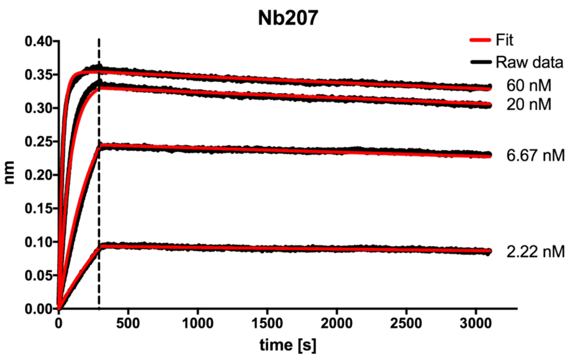

b

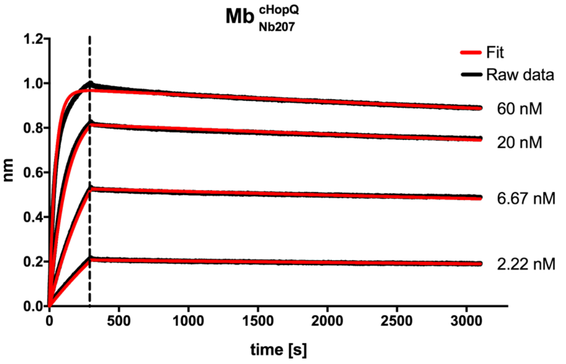

c

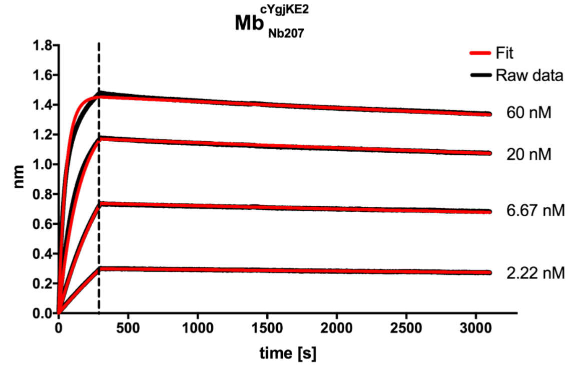

d

| | $K_D$ (pM) | $k_{on}$ x $10^5$ ($M^{-1}s^{-1}$) | $k_{off}$ x $10^{-5}$ ($s^{-1}$) |
|---|---|---|---|
| Nb207 | 38.2 | 6.530±0.015 | 2.51±0.01 |
| $Mb^{cHopQ}_{Nb207}$ | 76.8 | 4.018±0.008 | 3.09±0.01 |
| $Mb^{cYgjKE2}_{Nb207}$ | 86.8 | 3.346±0.007 | 2.90±0.01 |

**Supplementary Figure 6. Nb207, $Mb^{cHopQ}_{Nb207}$ and $Mb^{cYgjKE2}_{Nb207}$ bind to the cognate antigen with similar affinities.** Sensograms of the association and dissociation of Nb207 (**a**), $Mb^{cHopQ}_{Nb207}$ (**b**) and $Mb^{cYgjKE2}_{Nb207}$(**c**) onto immobilized GFP. Biotinylated GFP was immobilized on a Streptavidin (SA) bio-sensor and the binding kinetics were monitored by bio-layer interferometry (BLI) on OctetRED96 (ForteBio). The measured responses (black lines) were fitted to a monophasic 1:1 binding model (red lines). **d,** Calculated kinetic parameters are shown as mean standard error of the mean (s.e.m.) from n = 3 independent experiments.

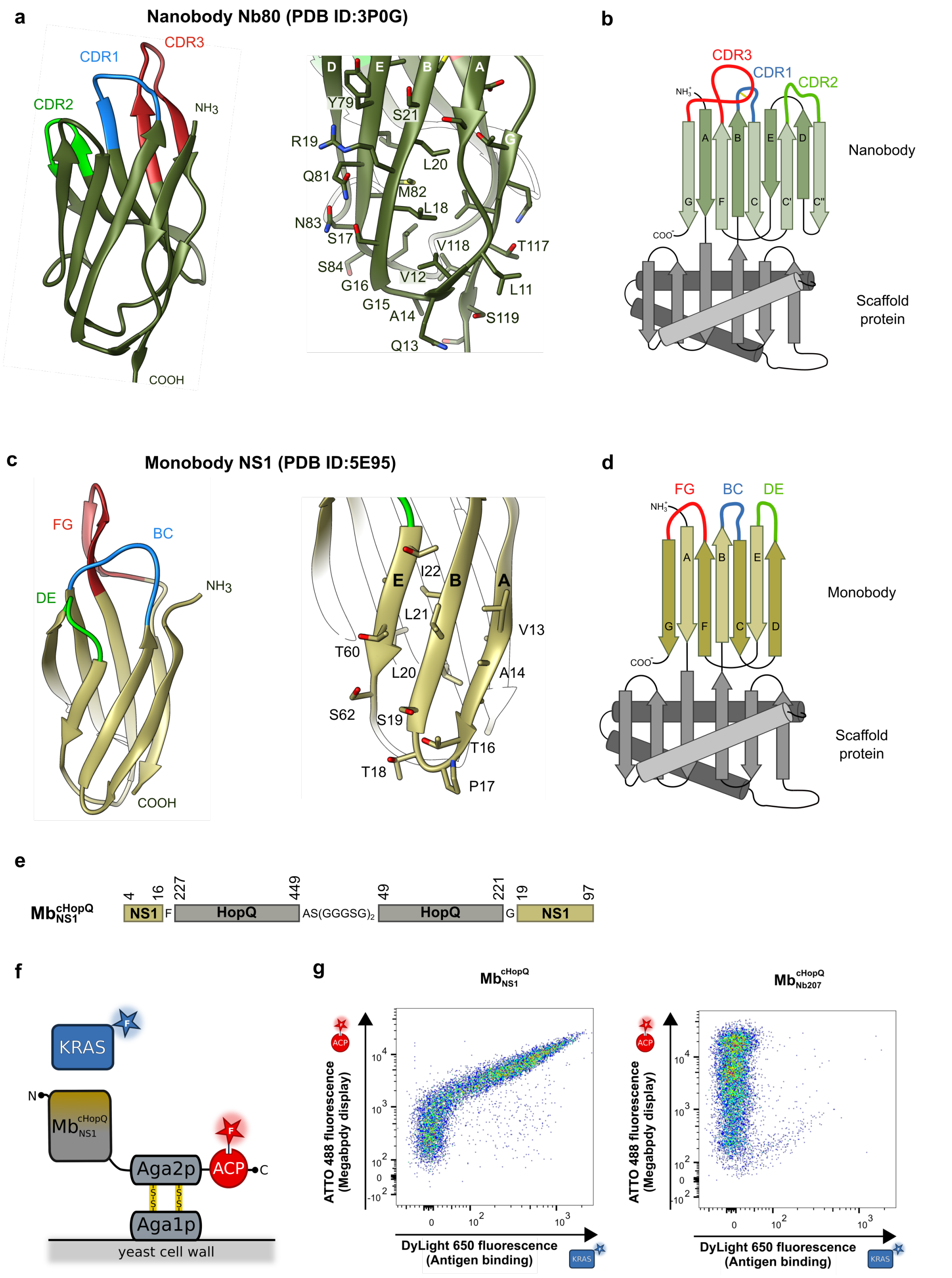

**Supplementary Figure 7. Yeast surface display of megabody $\text{Mb}^{\text{cHopQ}}_{\text{NS1}}$ built from the NS1 monobody that was grafted onto cHopQ. a,** Tertiary structure of a nanobody, based on crystal structure of Nb80 nanobody (PDB ID: 3P0G). The side chain conformations of β-strand A and β-strand B are indicated. **b**, Molecular design of a megabody that is assembled from a nanobody and a scaffold protein. **c**, Crystal structure of monobody NS1 (PDB ID: 5E95). The side chain conformations of β-strand A and β-strand B are shown. **d**, Molecular design of a megabody that is assembled from a monobody and a scaffold protein. **e**, Schematic representation of the primary structure of $\text{Mb}^{\text{cHopQ}}_{\text{NS1}}$. **f**, $\text{Mb}^{\text{cHopQ}}_{\text{NS1}}$ was displayed on the surface of yeast as a $\text{Mb}^{\text{cHopQ}}_{\text{NS1}}$-Aga2p-ACP fusion, and orthogonally stained with CoA-488 (red star) to monitor the display level. Binding of the antigen was monitored by incubating the yeast cells with 100 µM KRAS-DyLight 650 (blue star). **g**. Comparison of flow cytometric dotplots representing yeast cells displaying $\text{Mb}^{\text{cHopQ}}_{\text{NS1}}$ (left panel) and cells displaying $\text{Mb}^{\text{cHopQ}}_{\text{Nb207}}$ (right panel).

**a**

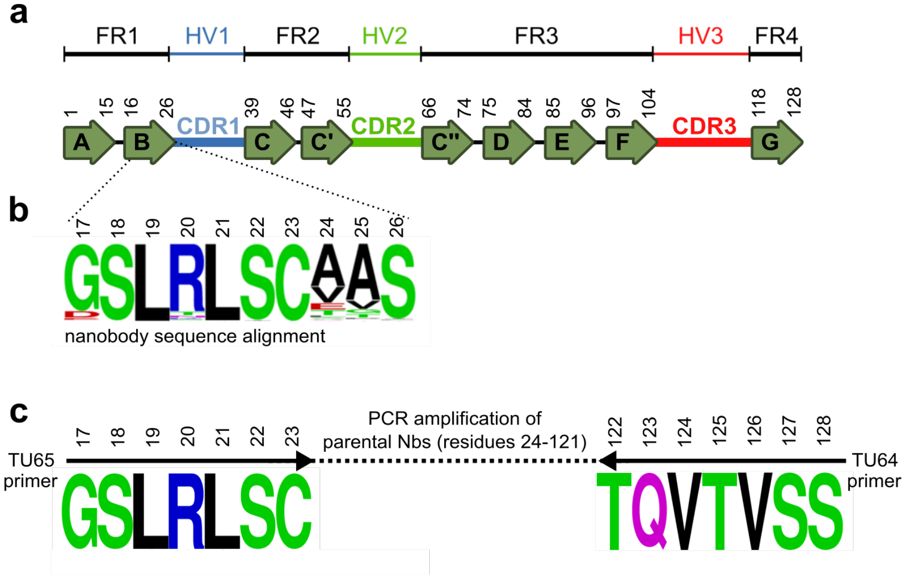

**d**

| selected clone | CDR1 | CDR2 | CDR3 |
|---|---|---|---|
| **MP1251_D10*** | GRTFSTAA | IYWTVGST | AARRRGFTLA PTRANE |
| **MP1251_D4** | GRTFSMSA | ITWSVGSP | AARRAGYVRL PTREDE |
| **MP1251_H6** | GGTFSISA | ISWNVGRP | AGRRAGYFGM PTREYE |
| **MP1251_A10** | GRTFSISL | IGWSTGNT | AARRRGWGPI PTMENE |
| **MP1251_A7** | GRTFSVSN | IGWTTGST | AARRRGYSRV PMTPDE |
| **MP1251_A5** | GRTSSNNH | ISWSRDFT | AAAPWGTNWR VKESE- |
| **MP1252_D10** | GRTFSLSM | ISWSRGST | AARRRGYISI PTTPDE |
| **MP1252_C10** | GRTFSMSA | ISWTVGRP | AARRAGYSRV PTREYD |
| **MP1263_C9** | GRTFSSDA | ISWSNAIT | AGGLNSYGST WAPQS- |

**e**

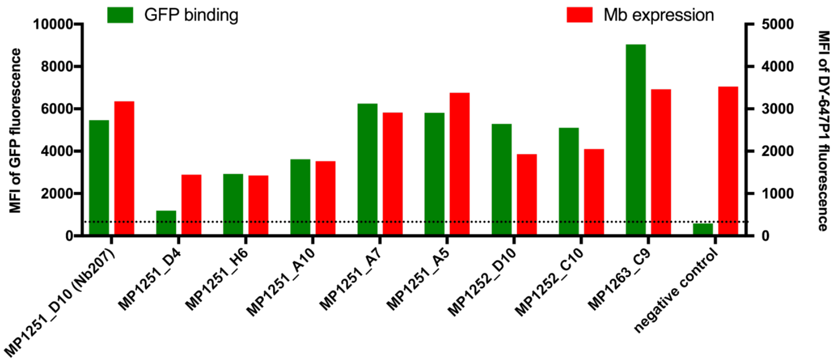

**Supplementary Figure 8. Sequences and binding properties of a representative set of GFP-specific megabodies selected by yeast-display a**, Schematic representation of a rearranged gene encoding a VHH domain (nanobody) in camelids. Conserved framework (FR, black) and hypervariable (HR, blue, green and red) regions are indicated and encode nine β-strand and three CDR regions, respectively. CDRs and β-strands of nanobodies are defined according to IMGT numbering. **b,** Alignment of β-strand B sequence, originated from 600 nanobody sequences available in-house (three different animals). **c**, PCR product of the *in vivo* maturated nanobody immune libraries amplified using TU65 and TU64 primers (**Supplementary Table 3**). **b,** CDRs composition of the nine megabodies selected by yeast display. CDRs are defined according to IMGT. Selected megabody clone MP12551_D10 contains the same CDRs composition as the nanobody Nb207, which was discovered by phage display (data not shown). **c,** Flow cytometric analysis of GFP binding for i yeast clones displaying nine selected megabodies. Individual yeast clones were orthogonally staining with Co-647 and incubated with 100 nM GFP. For each clone, the mean fluorescent intensities (MFI) of the DY-647P1 fluorescence (display level, red bars) and the GFP fluorescence (antigen binding, green bars) were calculated using the FlowJo software and compared to a cell displaying $Mb^{cHopQ}_{MP1031\_F2}$(the nanobody MP1031_F2 binds human coagulation Factor IX[8], negative control). The MFI of GFP fluorescence of a negative control is indicated as a dotted line.

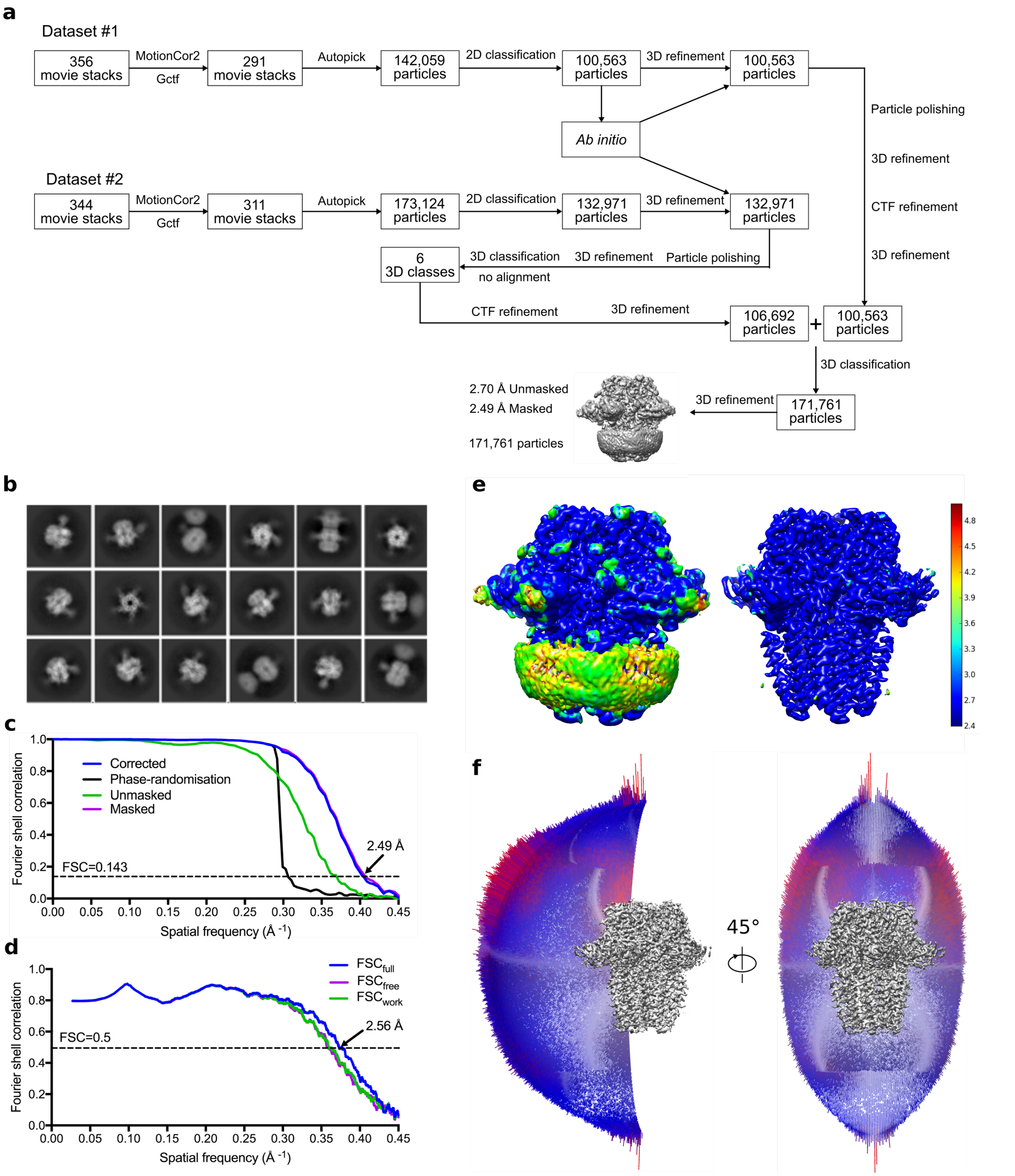

**Supplementary Figure 9. Cryo-EM image processing procedure for high resolution reconstruction of β3 GABA$_A$R in complex with to Mb$^{c7HopQ}_{Nb25}$. a.** Graphical overview of cryo-EM data collection and image processing (see Methods). **b,** 2D class averages used for cryo-EM map reconstructions. Aligned micrographs were obtained using FEI Titan Krios, Falcon3 detector and VPP (box size of 256 Å). **c,** FSC curves for the 3D reconstruction using gold-standard refinement in RELION. Data is shown for the phase randomisation, unmasked, masked and phase-randomisation-corrected masked maps. **d,** FSC curves for the atomic model refinement. Data is shown for model versus summed map (FSC$_{full}$), model refined in half-map 1 versus half-map 1 (FSC$_{work}$), and model refined in half-map 1 versus half-map 2 (FSC$_{free}$). **e**, Unsharpened cryo-EM map colored by local resolution (estimated using ResMap) shown at a lower contour level (left) and at a higher level (right). **f**, Angular-distribution histogram of particles used in calculating the final 3D reconstruction for the of histamine bound β3 GABA$_A$ receptor in a complex with Mb$^{c7HopQ}_{Nb25}$.

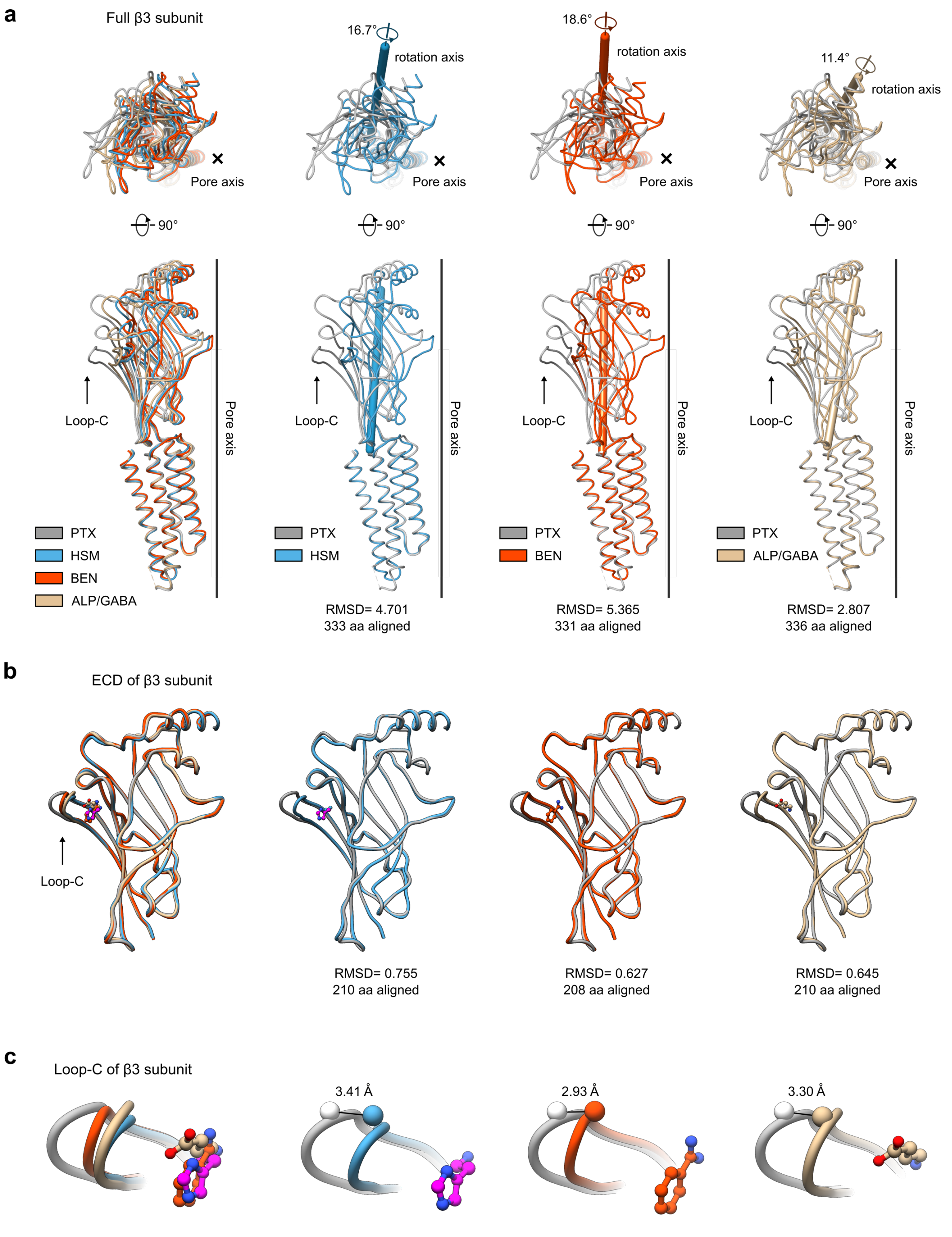

**Supplementary Figure 10. Structural analysis of β3 subunits of PTX-bound, ALP/GABA-bound α1β3γ2L $GABA_A$ receptor and BEN-bound, HSM-bound β3 $GABA_A$ receptor structures. a-c** Superposition of full β3 subunits (**a**), ECD (**b**) and Loop-C (**c**) of PTX-bound α1β3γ2L $GABA_A$ (grey, PDB ID: 6HUG), ALP/GABA-bound α1β3γ2L $GABA_A$ (khaki, PDB ID: 6HUO), BEN-bound β3 $GABA_A$ (orange, PDB ID: 4COF) and HSM-bound β3 $GABA_A$ (blue, PDB ID: 6QFA) receptors. **a** Superposition of full subunits on the basis of the global TMD alignment reveals the relative β3 ECD motions upon binding to PTX, ALP/GABA, BEN and HSM where rotation axis (sticks) and angles are indicated. The RMSD values are shown for full β3 subunits. **b** Superposition of ECDs (residues 1-217), where RMSD values are shown for ECDs. **c** Superposition of Loop-C, where differences in distances (Å) between the selected Thr202 $C_\alpha$ atoms (shown as spheres) are indicated with lines. The bound histamine (HSM), benzamidine (BEN) and GABA are indicated in magenta, orange and khaki, respectively.

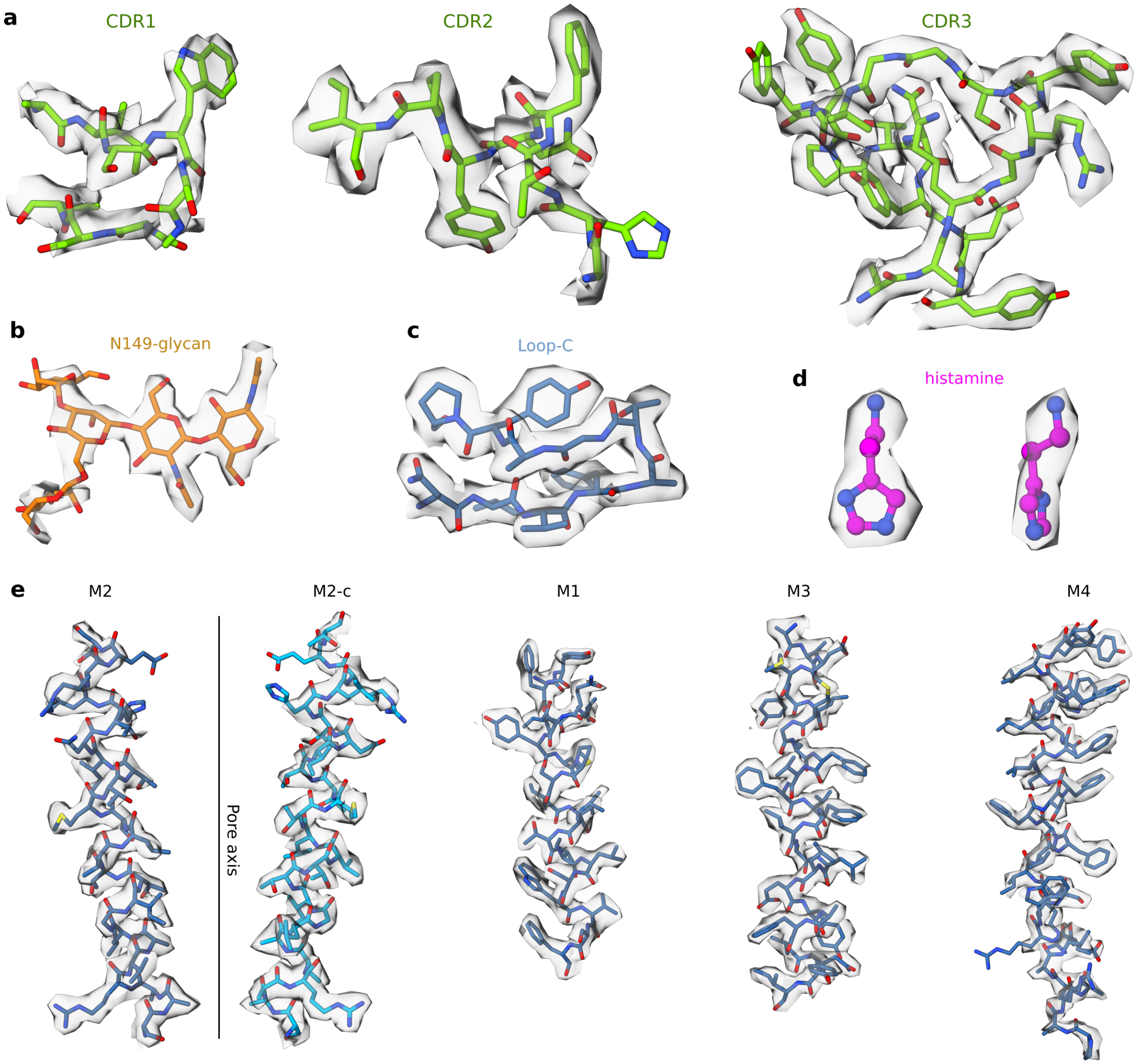

**Supplementary Figure 11. Histamine-bound β3 $GABA_A$ receptor model-map validation and electron microscopy density. a-e,** Electron microscopy density segments of $Mb^{c7HopQ}_{Nb25}$ CDRs (**a**) and β3 $GABA_A$ receptor N149-glycan (**b**), Loop-C (**c**), histamine (**d**), α-helices of TMD regions (**e**) (EMDB ID: 4542, PDB ID: 6QFA). Sharpened density maps are contoured at 0.08.

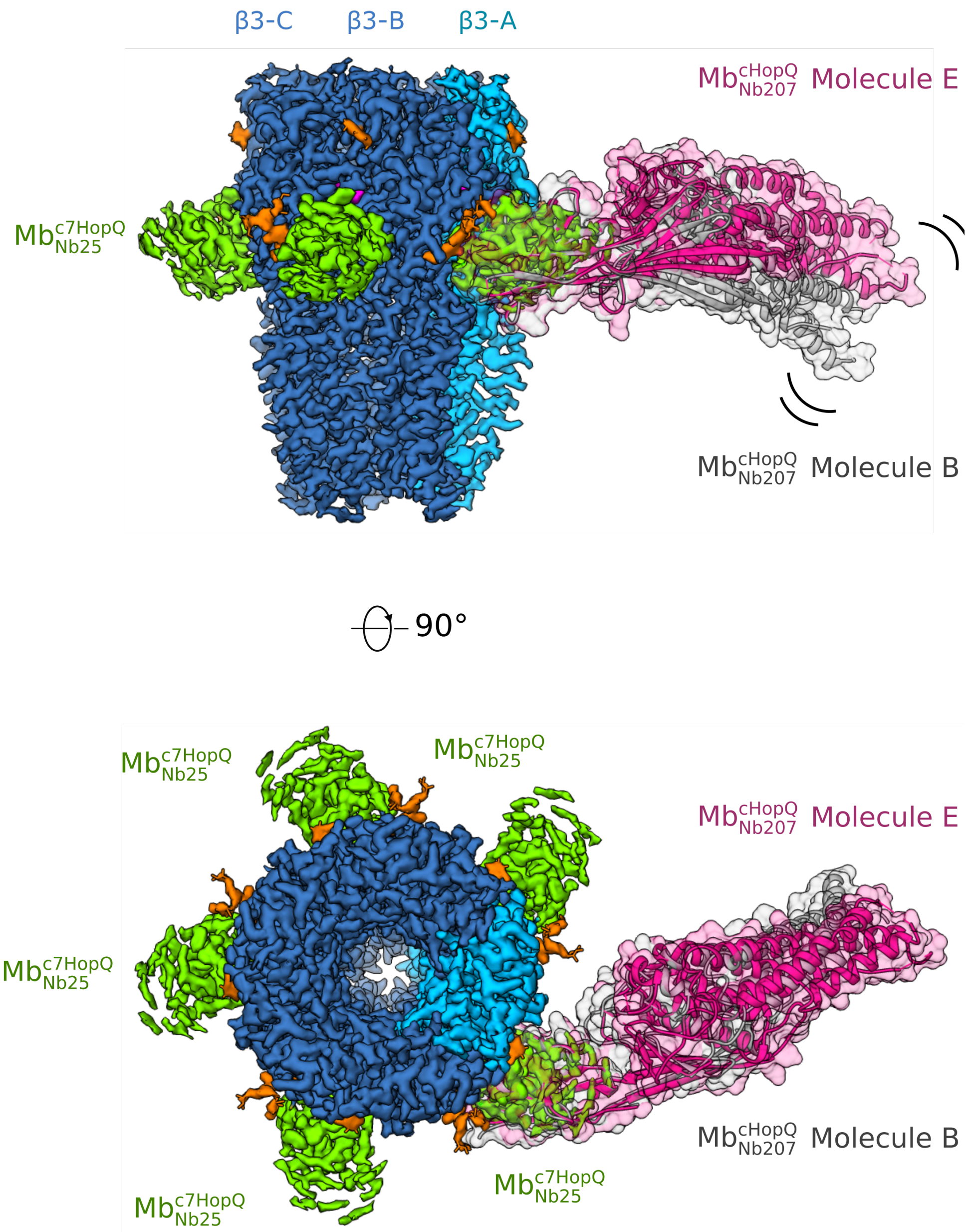

**Supplementary Figure 12. Molecular docking of the $\mathbf{Mb^{cHopQ}_{Nb207}}$ crystal structure onto the cryoEM map of the β3 GABA$_A$ receptor in a complex with $\mathbf{Mb^{c7HopQ}_{Nb25}}$.** The two most distinct $Mb^{cHopQ}_{Nb207}$ molecules from the asymmetric unit of the crystal structure (molecules B and E, PDB ID: 6QD6) are coloured in grey and magenta, respectively. They were aligned to the part of $Mb^{c7HopQ}_{Nb25}$ that was refined in the cryo-EM structure of the β3 GABA$_A$R in a complex with $Mb^{c7HopQ}_{Nb25}$ (EMDB ID: 4542, PDB ID: 6QFA).

**Supplementary Table 1**. Amino acid sequence of Nb207.

| **Nb207** |
| --- |
| QVQLQESGGGLVQAGGSLRLSCAASGRTFSTAAMGWFRQAPGKERDFVAGIYWTVGSTY YADSAKGRFTISRDNAKNTVYLQMDSLKPEDTAVYYCAARRRGFTLAPTRANEYDYWG QGTQVTVSS |

**Supplementary Table 2**. Data collection and refinement statistics.

| | $Mb^{cHopQ}_{Nb207}$ |
| --- | --- |
| **Data collection** | |
| Space group | P1 |
| Cell dimensions | |
| *a, b, c* (Å) | 71.17, 92.92, 244.22 |
| α , β , γ (°) | 92.05, 96.93, 112.15 |
| Resolution (Å) | 41.45-2.84 (2.90-2.84) * |
| $R_{meas}$ | 0.05 (0.66) |
| $I / \sigma I$ | 11.65 (1.45) |
| Completeness (%) | 95.6 (94.5) |
| Redundancy | 1.78 (1.77) |
| **Refinement** | |
| Resolution (Å) | 38.85-2.84 (2.90-2.84) |
| No. reflections | 230255 (13649) |
| $R_{work} / R_{free}$ | 0.225/0.251 |
| No. Atoms | |
| Protein | 33077 |
| Ion $Cl^-$ | 1 |
| Water | 114 |
| B factor | |
| Protein | 99.9 |
| Ion $Cl^-$ | 91.9 |
| Water | 91.7 |
| R.m.s. deviations | |
| Bond lenghts (Å) | 0.02 |
| Bond angles (°) | 1.87 |

*Values in parentheses are for highest resolution shell.

**Supplementary Table 3.** Cryo-EM data collection, refinement and validation statistics.

| | β3 $GABA_AR$ - $Mb^{c7HopQ}_{Nb25}$ complex<br>EMDB: 4542<br>PDB: 6QFA |
|---|---|
| **Data collection and processing** | |
| Microscope, location | Krios-II, MRC-LMB |
| Magnification | 75,000 |
| Voltage (kV) | 300 |
| Detector | Falcon 3EC with VPP |
| Electron Dose ($e^-$/Å$^2$) | 30 |
| Exposure time (s) | 60 |
| Pixel Size (Å) | 1.07 |
| Dose rate ($e^-$/pixel/s) | 0.4 |
| Frame number | 75 |
| Defocus Range (μm) | -0.7 to -0.5 |
| Micorgraphs collected (no.) | 700 |
| Micorgraphs selected (no.) | 602 |
| Initial particle images (no.) | 315,183 |
| Final particle images (no.) | 171,761 |
| Symmetry imposed | C5 |
| Map resolution (Å) | 2.49 |
| FSC threshold | 0.143 |
| Map resolution range (Å)[a] | 2.25-5.95 |
| **Refinement** | |
| Initial model used (PDB code) | 4COF, 5O8F |
| Model resolution (Å)[b] | 2.56 |
| FSC threshold | 0.5 |
| Model resolution range (Å) | 2.56 |
| Map sharpening *B* factor (Å$^2$) | -68 |
| Model composition | |
| Protein residues | 2,270 |
| Non-hydrogen atoms | 18,755 |
| Protein atoms | 18,240 |
| N-linked glycan atoms | 375 |
| HSM atoms | 40 |
| R.m.s. deviations | |
| Bond lengths (Å) | 0.005 |
| Bond angles (°) | 0.761 |
| Validation | |
| MolProbity score | 1.28 |
| Clashscore | 5.3 |
| Poor rotamers (%) | 0 |
| Ramachandran plot | |
| Favoured (%) | 98.17 |
| Allowed (%) | 1.83 |
| Disallowed (%) | 0 |

[a]Local resolution range. [b]Resolution at which FSC between map and model is 0.5.

**Supplementary Table 4**. Primer list.

| | **5' – 3' sequence** |
|---|---|
| EP230 | AGGACTGCTCTTCCACTGGAGACGGTGACCTGGGT |
| TU64 | CCCTCCACCAGAGCCACCTCCCAAGCTTGAGACGGTGACCTGGG |
| TU65 | GCATGTAACCACATCAAAGTATGGATCCCTGAGACTCTCCTG |
| TU89 | CCTTGAGCTCTTCGTCCCTGAGACTCTCCTG |
